# Tolerance to *Haemophilus influenzae* infection in human epithelial cells: insights from a primary cell-based model

**DOI:** 10.1101/2023.07.11.548529

**Authors:** Ulrike Kappler, Anna Henningham, Marufa Nasreen, Andrew H. Buultjens, Timothy P. Stinear, Peter Sly, Emmanuelle Fantino

**Affiliations:** School of Chemistry and Molecular Biosciences, The University of Queensland, St Lucia, Qld 4072, Australia; Child Health Research Centre, The University of Queensland, South Brisbane, Qld 4101, Australia; Department of Microbiology and Immunology, Doherty Institute for Infection and Immunity, University of Melbourne, Melbourne, Vic 3000, Australia

## Abstract

*Haemophilus influenzae* is a human respiratory pathogen and inhabits the human respiratory tract as its only niche. Despite this, the molecular mechanisms that allow *H. influenzae* to establish persistent infections of human epithelia are not well understood.

Here, we have investigated how *H. influenzae* adapts to the host environment and triggers the host immune response using a human primary cell-based infection model that closely resembles human nasal epithelia (NHNE).

Physiological assays combined with dualRNAseq revealed that NHNE from five healthy donors all responded to *H. influenzae* infection with an initial, ‘unproductive’ inflammatory response that included a strong hypoxia signature but did not produce pro-inflammatory cytokines. Subsequently, an apparent tolerance to large extra- and intracellular burdens of *H. influenzae* developed, with NHNE transcriptional profiles resembling the pre-infection state. This occurred in parallel with the development of intracellular bacterial populations, and appears to involve interruption of NFkB signalling. This is the first time that large-scale, persistence-promoting immunomodulatory effects of *H. influenzae* during infection have been u. Interestingly, NHNE were able to re-activate pro-inflammatory responses towards the end of the 14-day infection resulting in release of pro-inflammatory cytokines (IL8, TNFα). Our data further indicate the presence of infection stage-specific gene expression modules, highlighting fundamental similarities between immune responses in NHNE and canonical immune cells, which merit further investigation.

## Introduction

Chronic diseases of the respiratory tract, such as asthma, Chronic Obstructive Pulmonary Disease (COPD), and bronchiectasis, are highly prevalent, and infections with bacterial respiratory pathogens frequently affect these patients and hasten disease progression ^1–3^. Despite this, insights into the molecular events that occur at the interface between human epithelia and bacterial pathogens are rare at present.

*Haemophilus influenzae* is a human-adapted pathobiont that inhabits the nasopharynx as a commensal but causes disease in other parts of the respiratory tract ^1–3^. Currently, non-typeable strains of *H. influenzae* (NTHi) are the most common type of clinical isolate, and in addition to causing acute diseases such as otitis media and pneumonia ^2^, these strains are a major cause of exacerbations of chronic lung diseases, including in patients recovering from COVID-19 ^4–11^. While NTHi infections are rarely lethal, their high frequency and prevalence in patients suffering from chronic lung diseases make them a major driver of healthcare costs ^12, 13^. Additionally, NTHi infections affect particularly vulnerable groups, such as the young and the elderly, and are overrepresented in indigenous populations worldwide ^2, 14, 15^. Combined with the rise of resistance to beta-lactam antibiotics and the emergence of multidrug-resistant strains ^16, 17^, new insights into the biology of NTHi are urgently needed.

A key factor in NTHi virulence are interactions between the bacteria and the respiratory epithelia. Despite this, insights into the molecular interactions that allow NTHi to persist in contact with human epithelia are lacking, but likely hold the key to uncovering both bacterial and host processes that are crucial for infection.

NHBE or NHNE (normal human nasal epithelia) are fully-differentiated epithelia derived from primary human cells cultured at the air-liquid interface, and are an emerging model for bacterial infections of human respiratory epithelia ^18–23^. NHNE and NHBE contain basal, goblet and ciliated cells and accurately approximate human respiratory epithelia, the preferred niche of NTHi, including replicating authentic transcriptional responses to infection ^24^. Since their emergence, several studies have reported NTHi-NHNE infection data. However, the data were generally limited to short-term infections (up to 72h) and assessed changing bacterial burdens, but have provided little data tracking the molecular interactions between pathogen and host cells ^18, 20, 21, 25, 26^.

Only a single study that used NHBE derived from a single human donor has reported molecular insights into NHBE-NTHi interactions ^23^. The infection with the otitis media isolate strain, NTHi176, lasted 72h, at which point significant damage to the NHBE was observed ^23^. Dual RNAseq revealed complex adaptive processes in both the host and bacteria. NTHi infections reduced expression of NHBE genes encoding cytoskeleton elements and junctional complexes. In contrast, genes involved in pathogen recognition (ICAM1, SPON2, CEACAM7), extracellular matrix formation and pro-inflammatory cytokines (IL8, IL1β, CXCL5, 9, 10 & 11) were upregulated ^23^. Simultaneously, in NTHi 176, virulence factors such as key adhesins, amino acid biosynthesis pathways, several transport proteins required for the uptake of nutrients and known stress response proteins were upregulated ^23^.

To close the gap in our understanding of long-term colonization of respiratory epithelia by NTHi in human populations, we have used primary human nasal epithelia (NHNE) derived from 5 healthy donors and monitored NTHi infections over 14 days. Physiological and molecular data for the host and bacteria were collected prior to infection and 1-, 3-, 7-, and 14-days post-infection (p.i.), and revealed that while NTHi gene expression profiles underwent a major transition in the first 24 h of infection, subsequent changes were comparatively subtle. In contrast, NHNE transitioned through a strong hypoxia-driven response on day1 p.i. to a state resembling the pre-infection gene expression profile by day3 p.i., despite increasing NTHi loads. In keeping with this tolerance of NTHi infections, production of pro-inflammatory cytokines was delayed and only showed appreciable increases from about day7 p.i., coinciding with an increase in host cell death signalling and stress responses in NTHi.

## Results

### Long-term colonization of differentiated normal human nasal epithelia (NHNE) by non-typeable *Haemophilus influenzae* (NTHi) has minimal effects on epithelial integrity or health

A hallmark of NTHi infections in humans is their high level of persistence, both during ‘carriage’ of NTHi strains as nasopharyngeal commensals and during infections of other areas of the respiratory tract ^27, 28^. To investigate the dynamics of long-term interactions between NTHi and human respiratory epithelia, NHNE derived from five healthy human donors were infected with *H. influenzae* 86-028NP, and stable co-cultures with both extra- and intracellular NTHi populations developed within 24 h of infection (Figure 1, Figure S1). Over the course of the 14-day infection, total NTHi cell numbers increased slightly from 4.28 × 10^7^ average CFU/ml on day1 (24h) p.i. to 6.22 × 10^7^ average CFU/ml by day14. Within the same time period, intracellular populations of NTHi increased approximately 10-fold from 2.4 × 10^5^ CFU/ml or 0.5% of total NTHi present on day1 (24 h) p.i. to a maximum of 2.76 × 10^6^ CFU/ml or 4.4% of total NTHi present on day14. Particularly the development of intracellular NTHi populations showed donor-specific differences, increasing most slowly for donor 2 and fastest for donor 4 (Figure S1).

**Figure 1.**
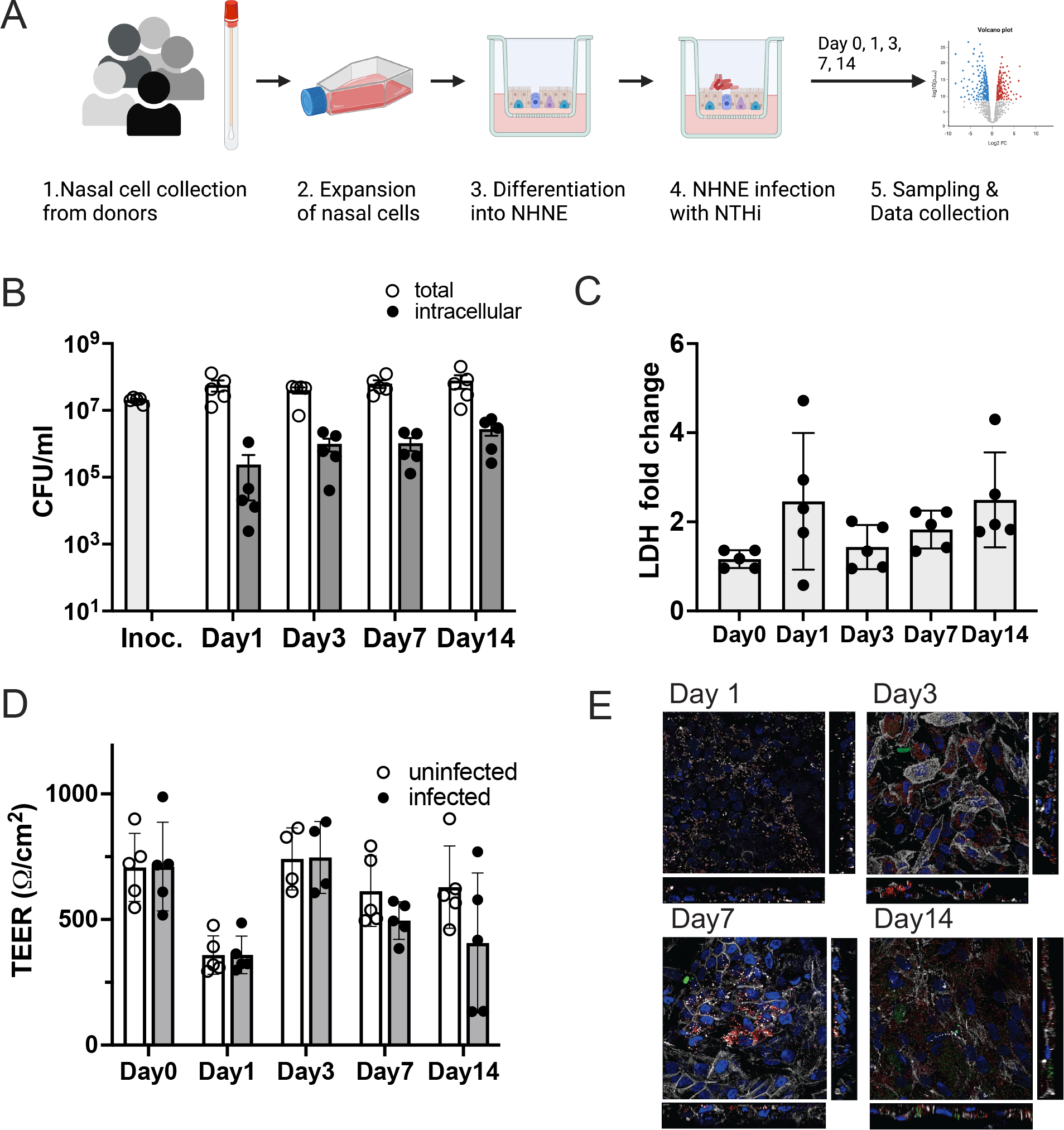
Infection of NHNE from five healthy human donors with *H. influenzae* strain 86-028NP Panel A: Schematic representation of experimental workflow. Prepared with BioRender.com. **Panel B: Bacterial loads during infection.** Total bacterial and intracellular bacterial loads (CFU/mL) were determined on Day 1, Day 3, Day 7 and Day 14. Each dot represents the average CFU/mL observed for each donor (*n*=2 biological replicates for each donor and condition). Bars and error bars indicate average CFU/mL and standard error. Inoc is the CFU/mL in the bacterial inoculum used to infect NHNE. **Panel C: Changes in LDH-release in infected NHNE.** LDH levels in infected and uninfected samples were determined using AlphaLISA. The fold change in infected compared to uninfected NHNE from each donor is shown, averages and standard error are shown. **Panel D: Transepithelial resistance (TEER) changes resulting from NHNE infection**. TEER readings for uninfected and infected NHNE for all donors are shown. Due to instrument failure, no readings were available for Donor 1 samples on Day 3 of infection. Average reading and standard error are shown. **Panel E: Immunofluorescence Microscopy of infected NHNE.** Representative images show donor 3 samples. White – Phalloidin stand of filamentous actin; Blue – cellular nuclei, DAPI stain; Green – TUNEL labelling; Red – anti-NTHi antibody. In Panels A, B &C, data plotted for each timepoint represent the average of multiple biological replicate measurements for samples from an individual donor.

These significant bacterial loads only caused minimal disruption of NHNE epithelial integrity, as shown by transepithelial electrical resistance (TEER) measurements that were consistent across infected and uninfected NHNE (Figure 1B). The only exception were the infected day14 samples from donors 4 & 5, where a decrease in TEER indicated epithelial damage. Lactate Dehydrogenase (LDH) release from NHNE as a measure of NTHi-induced damage and cell death confirmed these results. LDH levels only increased by 2-4 fold, with the highest levels detected during a transient increase on day1 p.i. (Figure 1C). During the infection, NTHi were present on the epithelial surface, inside the epithelial structure and close to the basal membrane and appeared more concentrated in specific areas of the epithelia (Figure 1D). However, at the sampling times used in this experiment, a preferential association of NTHi with ciliated cells, as reported by ^23^ for infection time points in the first 24h p.i., was not apparent.

### NTHi-infection of NHNE leads to dynamic, adaptive transcriptome changes

Although the physical infection parameters suggested a mostly stable coexistence of NTHi and NHNE, evaluation of transcriptional responses revealed highly dynamic, adaptive processes taking place during infection. To capture the adaptive changes occurring over the course of the infection, data were analyzed relative to the previous time point.

For NTHi, significant changes in gene expression were only observed during transition to the host environment in the first 24 h of infection, with all p.i.-time points clustering together in PCA plots (Figure 2A). In keeping with this, statistically significant changes in gene expression (2-fold, FDR p-value <0.05) were only observed for the Day1-Day0 comparison and 21 NTHi genes (Figure 2A, Table S1). Adaptation of NTHi to the host environment was primarily metabolic, with 11 of the 21 differentially expressed genes (DEGs) encoding enzymes required for *de novo* biosynthesis of purines, a uracil permease and an enzyme needed for the production of queuosine, a guanine derivative involved in tRNA modifications ^29^. Two enzymes associated with the partial NTHi TCA cycle, malic enzyme that interconverts pyruvate and malate and an NADP-specific glutamate dehydrogenase that produces alpha-ketoglutarate from glutamate, were also upregulated, as was the GlpX Class II bisphosphofructokinase which could indicate an increase in gluconeogenesis.

**Figure 2.**
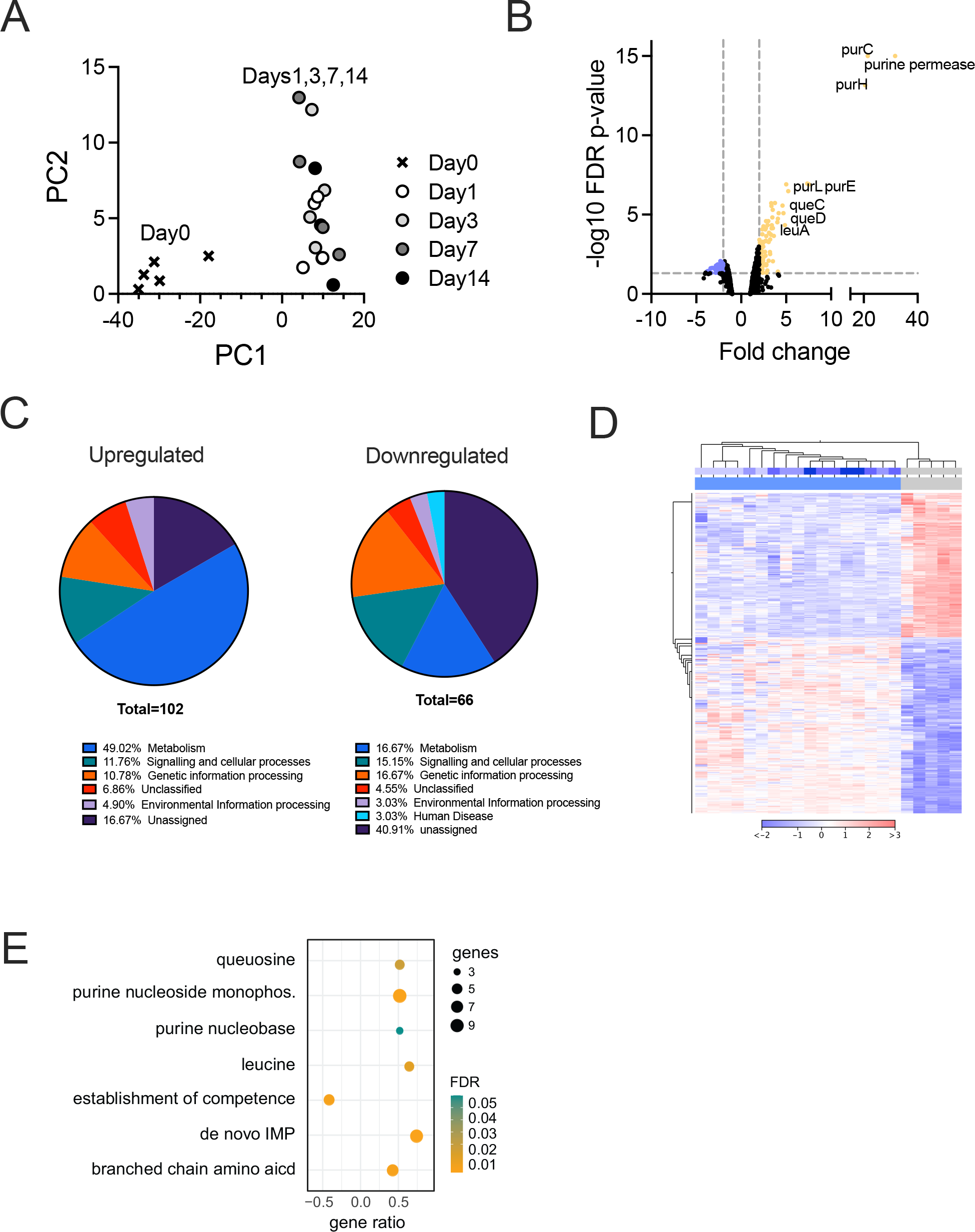
Remodelling of the NTHi 86-028NP transcriptome during NHNE infection. **Panel A: PCA plot** for NTHi whole transcriptome data. **Panel B: Volcano plot** of Differentially Expressed Genes (DEGs) for comparison of the combined day1-day14 samples against the pre-infection state (‘day0’). Colours indicate genes with at least a 2-fold change in expression (down-blue, up-yellow) and an FDR p-value <0.05. **Panel C: Distribution of NTHi DEGs into functional categories** using KEGG Orthology. Results were generated using BlastKOALA. **Panel D: Heatmap and dendrogram for NTHi DEGs (2-fold change, FDR p-value <0.05).** Metadata bars at the top of the heatmap: Lower bar – infected (blue) and uninfected (grey) samples, top bar – time post-infection, day0 / uninfected - grey, day1 to day14 – increasingly deeper shades of blue. **Panel E: GO-term enrichment for NTHi DEGs** identified in the uninfected vs infected (all) comparison. The GO term enrichment was generated using geneontology.org.

Upregulation of the MsgA methylglyoxal synthase suggests a role for conversion of glycolytic intermediates via methylglyoxal to lactate and pyruvate during NTHi host contact. Lastly, catalase upregulation (2.7-fold) indicates an increase in oxidative stress during the transition to the host environment, although catalase was the only gene of the NTHi OxyR oxidative stress regulon identified here ^30^.

### Global analysis confirms that major NTHi gene expression changes during infection are associated with nucleobase and amino acid metabolism

As the PCA plot revealed only modest transcriptional changes over time in NTHi samples, we also analyzed NTHi gene expression changes in pre-vs. post-infection samples (day0 vs. days1, 3, 7, 14). Here, 106 and 68 genes were up-or downregulated at least 2-fold with an FDR p-value <0.05 (Figure 2A-D, Table S1).

Among the 106 upregulated genes, metabolism (n=43) was the dominant category. This included purine biosynthesis (10 genes), queuosine biosynthesis (4 genes) and one gene in pyrimidine acquisition, confirming the results described above. Additionally, 11 genes involved in the synthesis of branched-chain amino acids that are essential in humans and thus would have limited bioavailability during infection were identified (Figure 2C, Table S1). In keeping with this, Gene Ontology (GO) analyses (biological process) revealed an overrepresentation of ‘IMP *de novo* biosynthetic process’ (20.7-fold), ‘purine nucleobase biosynthetic process’(17-fold), ‘queuosine biosynthetic process’ (14.2-fold) and ‘branched-chain amino acid metabolic process’ (12.2-fold) (Figure 2E, Table S1).

Upregulation of genes encoding transport proteins for purines, pyrimidines, queuosine, as well as peptides, amino acids and putrescine, underlines the limited availability of nucleobases and specific amino acids in the host environment (Table S1). Upregulation of stress responses included genes for resistance to reactive chlorine species and hydrogen peroxide (Table S1).

In contrast to the upregulated genes, most downregulated genes encoded hypothetical proteins or proteins of unknown function (26/68). Genes associated with competence and DNA uptake were significantly reduced in expression (31-fold enrichment in GO term analyses, Figure 2E), as was expression of genes encoding the alternative sigma factors RpoH and RpoE that, in other Gram-negative bacteria, regulate responses to cell envelope stress ^31^. Also noteworthy was the downregulation of the LctP lactate permease that is required for the uptake of L-lactate, a preferred carbon source for NTHi during infection ^19^, as well as downregulation of components of the TonB- and TolB-transport systems, a MexI-type RND efflux pump (also known as AcrB ^32^) and a Type II Secretion System (Table S1) that may have roles in virulence.

A global analysis of gene expression patterns revealed that for well over 50% of NTHi genes, expression levels decreased from day0 to day1 p.i. (Figure S2). We attribute this to the transition from *in vitro* growth on Brain-Heart Infusion, a nutrient-rich complex medium, to growth in the host environment where all nutrients must be obtained from the host cells or produced endogenously.

Genes encoding known NTHi virulence factors, reviewed, e.g. in ^1^, showed mixed expression patterns. While the opacity-associated protein, the HMW outer membrane proteins and ampicillin resistance were downregulated, the eP4 lipoprotein that is involved in NAD metabolism, the PntAB transhydrogenase and the peptidoglycan-associated OMP P6 (also known as Pal) showed increased expression on day1 p.i., and two genes encoding ferritin-like proteins (ftnA, and ftnA1) showed increased expression throughout, indicating an increased need to store and sequester iron in the host environment.

### Adaptation of NHNE to NTHi infection was dominated by a strong hypoxia response and activation of TLRs 4 & 9 on day1 p.i

The response of NHNE to infection differed markedly from that of NTHi, with significant and distinct changes in gene expression occurring throughout the 14-day infection (Figure 3A). In NHNE, the most significant gene expression changes occurred in the first 72 h p.i., where 884 (Day1-Day0) and 587 (Day14-Day7) genes were differentially up-or downregulated (>2-fold change, Bonferroni p-value <0.05), respectively. For the later stages of the infection process, 128 (Day7-Day3) and 139 (Day14-Day7) genes were differentially expressed (Figure 3A, B; Table S2).

**Figure 3.**
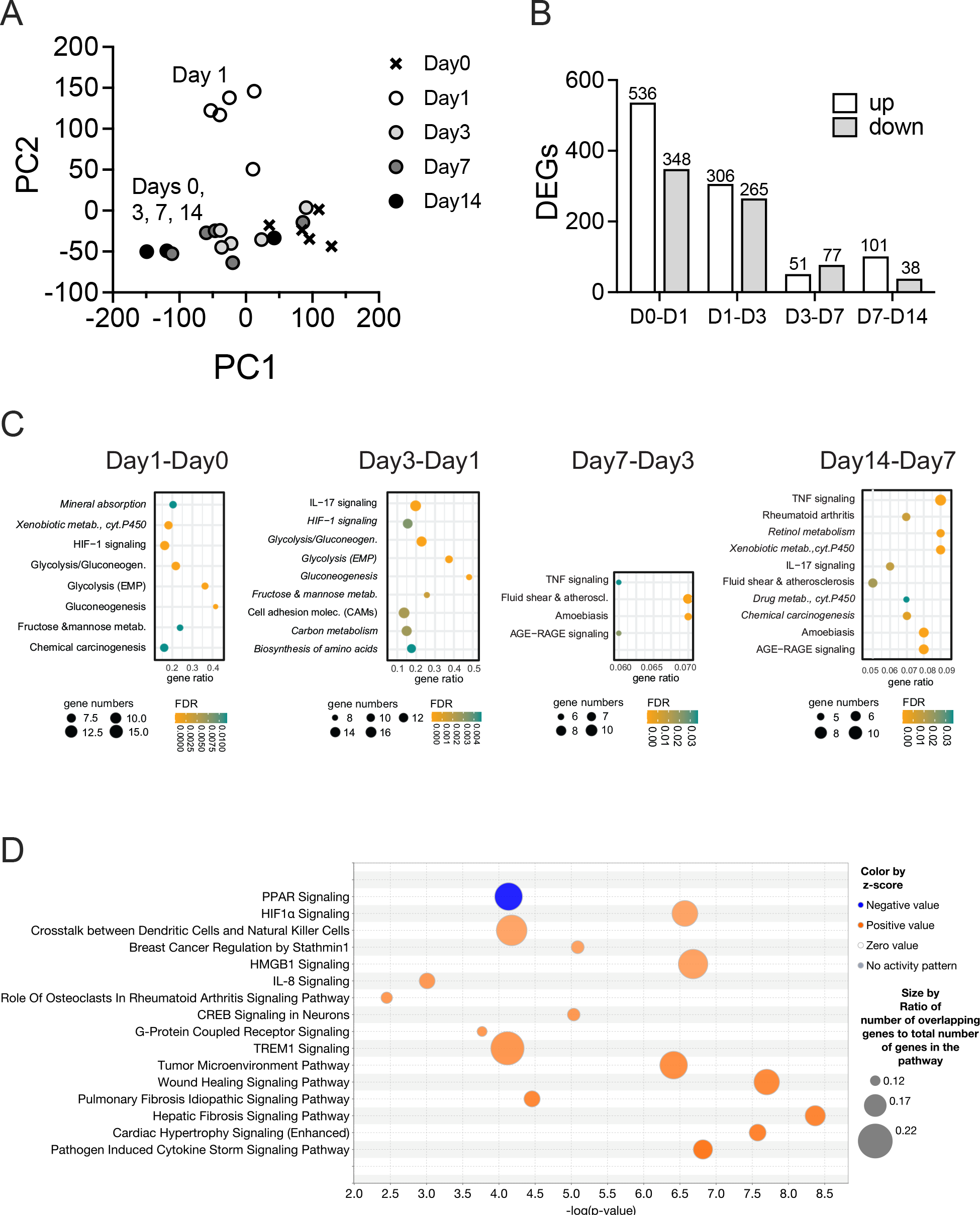
NHNE responses to NTHi infections. **Panel A:** PCA plot **Panel B**: Numbers of DEGs identified in NHNE over time. DEGs (≥2 fold expression change, Bonferroni p<0.05) were determined relative to the previous time point. **Panel C**: Gene Ontology analysis showing processes up-(regular font) and down-regulated (italics) in NHNE over time. Gene Ontology analyses used TOPPFUN, **Panel D**: Overrepresented pathways on day1 p.i. compared to day0 of NHNE infection. Pathways were identified using IPA (Qiagen) Canonical pathway analysis. Pathways with z-scores of 3.0 or higher and an FDR p-value <0.05 are shown. Pathways are displayed in ascending order of z-scores. Z-scores above 2.0 were considered significant.

On day1 p.i., NHNE gene expression changes were dominated by a strong upregulation of glycolysis-associated genes, glucose and lactate/pyruvate transporters (SLC2A1&3, SLC16A3) and lactate dehydrogenase (LDHA). This indicates a ‘Warburg-shift’ in NHNE metabolism that increases glycolytic flux, leads to lactate production from pyruvate and is associated with hypoxia signalling. Additionally, the antimicrobial peptide-producing ADM gene, several pro-inflammatory and pro-apoptotic factors (e.g. BNIP3, MIF, SERPINE1); PLAU, MMP9 and P4HA1 that modulate extracellular matrix composition and other known hypoxia-regulated genes such as carbonic anhydrase IX (CA9) were upregulated (Figure 4, Table S2). These gene expression changes were driven by HIF1α activation, identified by GO term analyses (TOPPFUN) (Figure 3C) and Ingenuity Pathway Analysis (IPA) that also revealed enrichment of pathways typically associated with bacterial infections such as ‘Pathogen-induced Cytokine Storm Signalling’, ‘Acute Phase Response Signalling’ and ‘Role of Pattern Recognition Receptors in Recognition of Bacteria and Viruses’. Underlying these higher-level, broad processes was activation of several more specific pathways such as IL6-, IL8-, TREM1- and CREB signalling (Figure 3D). The majority of all enriched pathways detected had functions in signalling, including immune cell attraction (Figure 3 C, D, Table S3).

**Figure 4.**
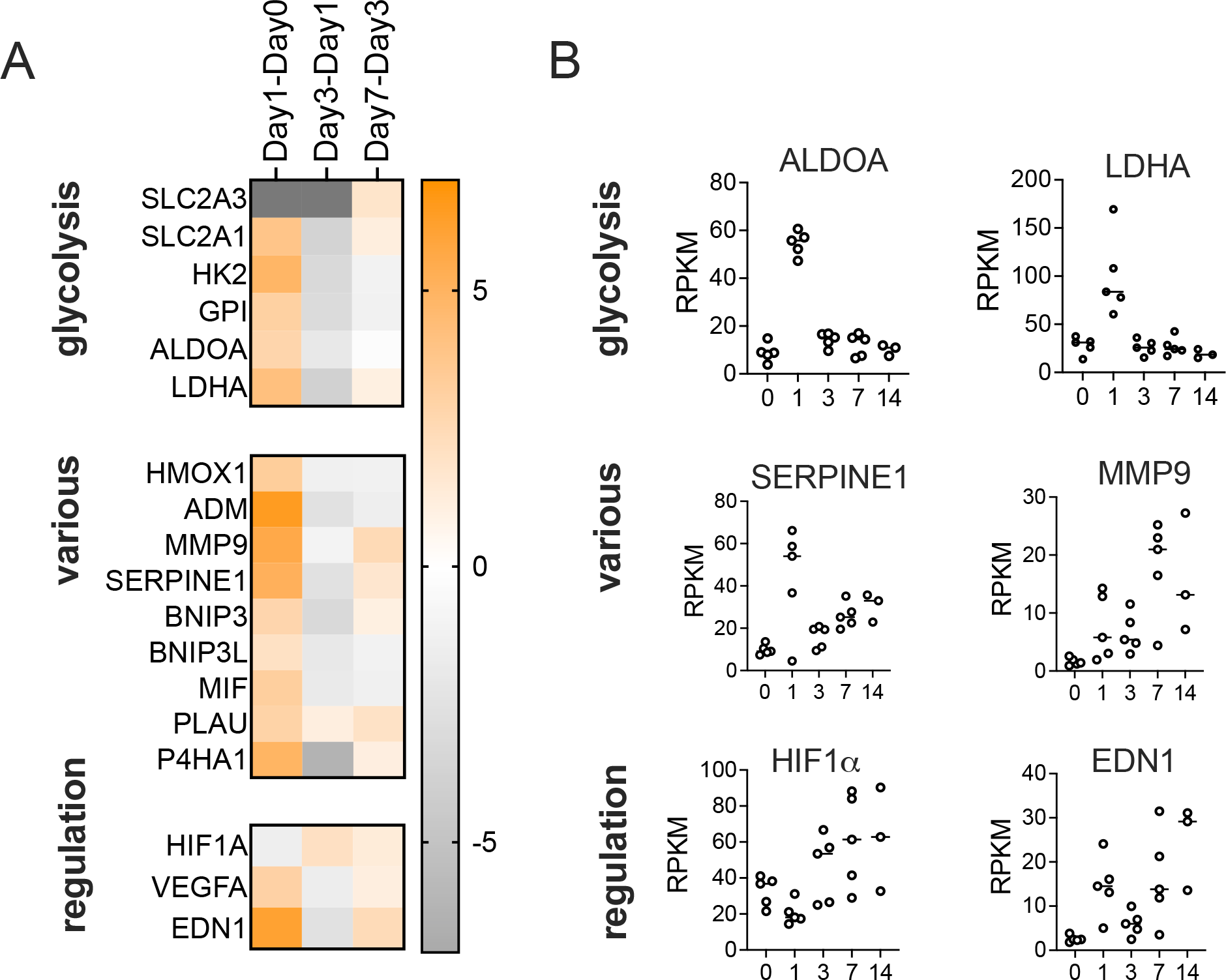
Hypoxia-driven NHNE response to infection on day1 p.i. **Panel A**: Heatmap showing fold changes in the expression genes involved in various relevant functions. Fold changes were determined using IPA canonical pathway analyses (Qiagen). **Panel B:** RPKM data for selected genes that show high fold-changes in expression on day1 compared to day0. Expression patterns of several regulators and effectors are complex and show clear changes both on day1 and later during the infection.

The activation of HIF1α was likely caused by exposure of NHNE to NTHi lipopolysaccharide (LPS), which was predicted to be the main effector driving changes in pathway and regulator activation at day1 p.i.. LPS-sensing is mediated by TLR4, and IPA analysis also suggests activation of TLR9 which, together with HIF1α activation led to increases in VEGFA, EDN1, and PLAUR levels and activation of a range of pro-inflammatory interleukins, including IL1A, IL1B, IL6, IL18 and TNFα, as revealed by upstream regulator analyses (IPA) (Figure S3). While overall low in expression, IL36A, which has been associated with lung infections ^33^, was also involved in this response (Figure S3).

The Day1-Day0 data analysis also revealed upregulation of the NUPR1 ferroptosis inhibitor and the alarmin HGMB1, which is involved in ECM remodelling via activation of matrix metalloproteases, was also observed. HGMB1 and TLR9 have both been previously identified as involved in otitis media caused by NTHi ^34^. The upregulation of TLR9 was likely related to the development of intracellular NTHi populations and the resulting presence of bacterial DNA in the cytoplasm of epithelial cells (Figure S3). Interestingly, at the same time as activating a pro-inflammatory response, the presence of NTHi also caused the activation of cellular damage repair pathways (hepatic fibrosis and wound healing pathways). This process requires increased lipid metabolism (GO:0006629, Biological Process, ToppFun) and, specifically, prostaglandin-related processes (GO:000669, Biological Process, ToppFun), both of which were overrepresented in GO-term analyses (Figures 3, S3).

### LPS-driven Hif1α and metabolic activation are reversed by day3 post-infection

Unexpectedly, at 72h p.i. (day3), NHNE gene expression patterns largely returned to a state similar to the uninfected state (day0), even though TNFα, IL1β and LPS remained relevant upstream regulators (Figures 3A, 4, S3). This included a reversal of the HIF1α-mediated hypoxia response and downregulation of genes associated with glycolysis and lactate production, suggesting a decrease in cellular glucose demand and possibly increased energy generation by respiration. In total, 316 genes (196 upregulated on day1, 120 downregulated on day1) underwent a reversal of gene expression compared to day1, returning mostly to pre-infection levels (Figure 3 A, B). This was also reflected in GO and pathway analyses, where HIF-1α signalling and glycolysis/gluconeogenesis pathways remained overrepresented but were downregulated (Figures 3C, 4, Table S3). Also downregulated were acute phase-, IL6-, IL8-, TREM1- and CREB-signalling and expression of TLR9 (Figure S3; Table S4).

The most highly upregulated genes in the Day3-Day1 comparison included genes of the canonical pathway ‘Pathogen-induced cytokine storm signalling’ and chemokines (CXCL5, CXCL9, CXCL10, CXCL11, CXCL14, CCL2), with both positive and negative regulatory functions in innate immune cell migration. This was reflected in the enrichment of canonical pathways such as IL-17 signalling, MAPK, & interferon signalling, and ‘Hypercytokinemia in Influenza Pathogenesis’ (Figure S3, Table S4).

### NTHi infection suppresses NHNE inflammatory responses

As a result of the continued upregulation of ‘Pathogen-induced cytokine storm signalling’ in NHNE expression data (Figure S3), we expected to see significant increases in the expression of genes encoding pro-inflammatory cytokines and interleukins. However, the corresponding pathways only showed positive enrichment on Day1-Day0, but not Day3-Day1.

This prompted us to further investigate the expression patterns of these genes and the presence of markers such as IL-8 and TNFα in basal media and NHNE apical wash fluids.

Interestingly, while the largest absolute fold-change in RPKM values occurred between day1 and day3 of infection, in absolute terms, gene expression levels were still modest on day3 and continued to rise until day14, with high levels only observed on days7 and 14 (Figure 5). In addition to TNFα, IL-8 and IL-1β, expression of Lipocalin2, which has proposed roles in sequestering iron during bacterial infection, prevention of apoptosis and cytokine secretion ^35, 36^ also strongly increased on days7 and 14.

**Figure 5.**
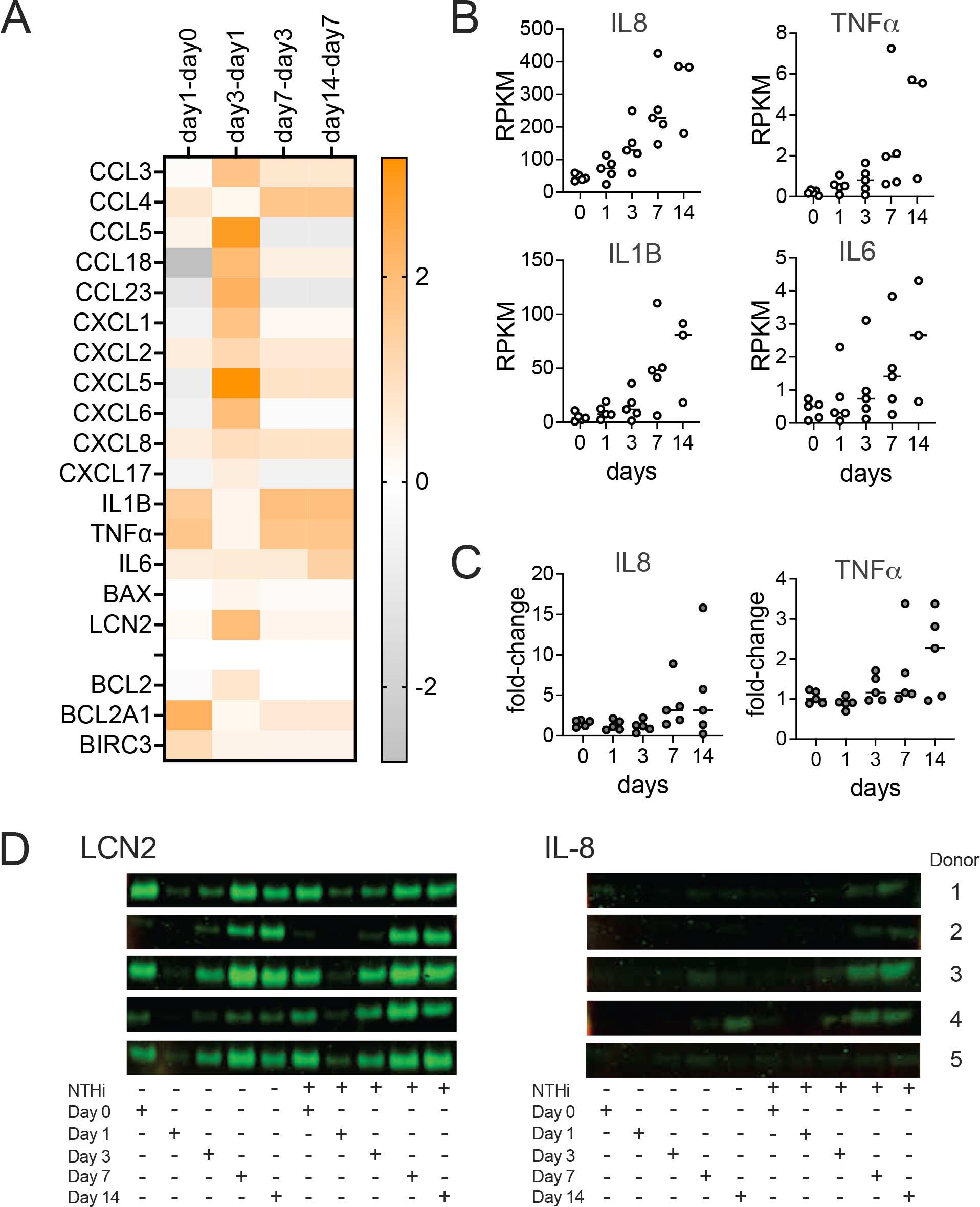
The NHNE immune response to infection is delayed following NTHi infection. **Panel A:** Heatmap showing fold-changes of genes involved in innate immunity over the course of a 14-day infection, **Panel B**: Gene expression changes over time for IL-8, TNFα, IL1β and IL-6. Data are shown as RPKM, **Panel C**: Fold-changes in the concentrations of IL-8 and TNFα in infected compared to uninfected NHNE. IL-8 and TNFα concentrations were detected using ELISA Kits and apical wash fluids. **Panel D**: Detection of IL-8 and LCN2(NGAL) in NHNE basal growth media by Western Blot. TNFα concentrations were below the detection limit.

To ascertain that the gene expression patterns reflect protein expression in our samples, levels of IL-8 and TNFα detected in apical washes and basal medium were determined and matched the gene expression data. Compared to uninfected NHNE, increases in IL-8 and TNFα in apical wash fluids by ELISA were modest. IL-8 levels increased less than 6-fold overall, except for Donor 2, where levels increased up to 15-fold on day14 p.i. Decreases in IL-8 levels observed on day14 for donors 4 & 5 are likely due to damage to the NHNE for these two donors that was also apparent in TEER measurements (Figures 1 & 5). TNFα levels were also very low throughout, with increases limited to a maximum of 3-4 fold on days 7 and 14 and highest increases observed for donor 4 & 5 NHNE that showed epithelial damage on day14 (Figure 5). Western Blot data for basal media samples confirmed that IL-8 was only detectable after day3 of infection (Figure 5 C, D), while TNFα was below the detection limit throughout.

High levels of Lipocalin2 were present in the apical wash fluids and the basal media at all timepoints, with a reduction in Lipocalin2 concentrations on days 1 and 3 p.i. for both uninfected and infected NHNE (Figure 5D, Figure S4). Together these data demonstrate that despite the early inflammation/ acute response signature apparent in the gene expression data, the pro-inflammatory response and production of signalling molecules was uncoupled from the metabolic and hypoxia-driven changes observed in the first 24-72 h of infection and significant production of these molecules delayed until lats stage infection (Figure 5). This has not previously been observed for NTHi infections of respiratory epithelia and suggests the existence of specific NTHi metabolites and proteins in this process.

### Apoptotic signalling and antimicrobial responses increased on days 7 and 14 post-infection

Compared to the significant changes in gene expression on day1 and day3 p.i., changes observed on day7 and day14 were less pronounced. Day7 gene expression patterns were enriched in canonical pathways associated with immune responses, such as ‘Macrophage Classical Activation Signalling’, IL17-, IL23- and pyroptosis signalling, and, at modest levels, HIF1α-, TREM1- and acute phase response signalling were also enriched (Figure S3; Table S4). The effectors controlling this were endothelin (EDN1), TGFB, the CCR5 chemokine receptor and two interleukins, IL17 and IL36A, that were also involved in the day1 response to NTHi infection (Figure S3). However, a strong hypoxia-related signature was missing on day7 (Figure 4), and several additional effectors that were upregulated on day1 were downregulated on day7 p.i., including IL10, which has anti-inflammatory properties, BCL6 that has been linked to susceptibility to influenza A infection, VIP, a major neuropeptide present in lung tissues and the LXR/RXR regulator pair ^37–41^.

The most highly upregulated gene on day7 was CYP1A1, a cytochrome P_450_, that is involved in xenobiotic metabolism and can be induced by activation of the aryl hydrocarbon receptor (AhR) ^42^. In macrophages, CYP1A1 can be induced by LPS exposure, where its expression continued to increase as infection progressed ^43^. During NTHi infection of NHNE, indole, produced by the bacteria via the reaction of the TnaA tryptophanase ^44^, could have led to activation of AhR ^45^.

Expression of the NTHi *tnaA* gene increased from day3 p.i., which matches the emergence of CYP1A1 as a highly expressed gene in the NHNE expression profile (Fig. S5).

Matching the data collected during the physical characterization of the NTHi-NHNE infection (Figures 1 & 5), expression levels of key chemo- and cytokines (e.g. CXCL8, IL6, IL1β, CCL3, CXCL1 & CXCL2) and genes involved in ECM remodelling and imflammation (PLAU, Serpine1, MMP9) increased further on day14. Expression of IER3 that protects cells from IL1β−& TNF-induced apoptosis, also increased in expression. Predicted active effectors on day14 included SOCS1 (suppressor of cytokine signalling 1), TRIM24, a co-activator of STAT3 expression, and ACKR2 that is involved in immune response silencing.

Both Day7-Day3 and the Day14-Day7 gene expression patterns showed increased TNFα- and AGE-RAGE signalling, indicating activation of NFkB and increased oxidative stress. This increase in innate immune signalling, inflammation and pyroptosis (Figure S3; Table S4) is likely related to the increase in apoptosis-associated DNA breaks observed by immunofluorescence microscopy of infected NHNE on days 7 and 14 (Figure 1D). While these changes were not significant in the global analyses, the increases in NHNE antibacterial processes on day7 and day14 p.i. were reflected in an increased expression of oxidative stress defence genes in NTHi. These included the peroxiredoxin encoding *pgdX* gene and genes involved in the SOS DNA repair response such as *lexA*, *recA*, *recN*, *uvrA*, *uvrD* and the NTHI_RS01685 single-stranded DNA-binding protein (Figure S5).

### Genes involved in cellular development and regulation of apoptosis may be increase susceptibility to NTHi-induced epithelial damage

Even though at a global level NHNE gene expression patterns showed no obvious differences for the five donors used in this study, NHNE derived from donors 4 & 5 showed clear signs of epithelial decay at day14 p.i. that were absent in donors 1-3. We hypothesized that this could be due underlying changes in gene expression and compared the two donor groups using both the DEG1 tool within the *iDEP* platform and univariate correlation, which identified 8 and 43 genes, respectively. Of the 43 genes identified by univariate correlation (abs. Pearson coefficient of > 0.75), many were expressed at very low levels, with 8 genes (TNFRSF12A, MAP7D1, ZNF107, CCDC88B, PHETA1, CD83, PHLDA2 and MRGPRX3) that had maximal RPKM values of >1 retained for further analysis (Figures 6, S6; Table S5). The most highly expressed gene in this group was TNFRSF12A, which has a role in the positive regulation of extrinsic apoptosis signalling and control of wound healing. TNFRSF12A forms a group with MAP7D1 that is required for autophagosome formation and CD83 and CCDC88B that are associated with regulating inflammation. MAP7D1 and CCDC88B are known to interact with cytoskeleton elements such as microtubules. The function of ZNF107 is not well defined other than that it is likely involved in gene regulation, and the remaining three genes have roles that are apparently unrelated to bacterial infection, namely in developmental processes (PHETA1, PHLDA2) and neuronal regulation and modulation of pain (MRGPRX3).

**Figure 6.**
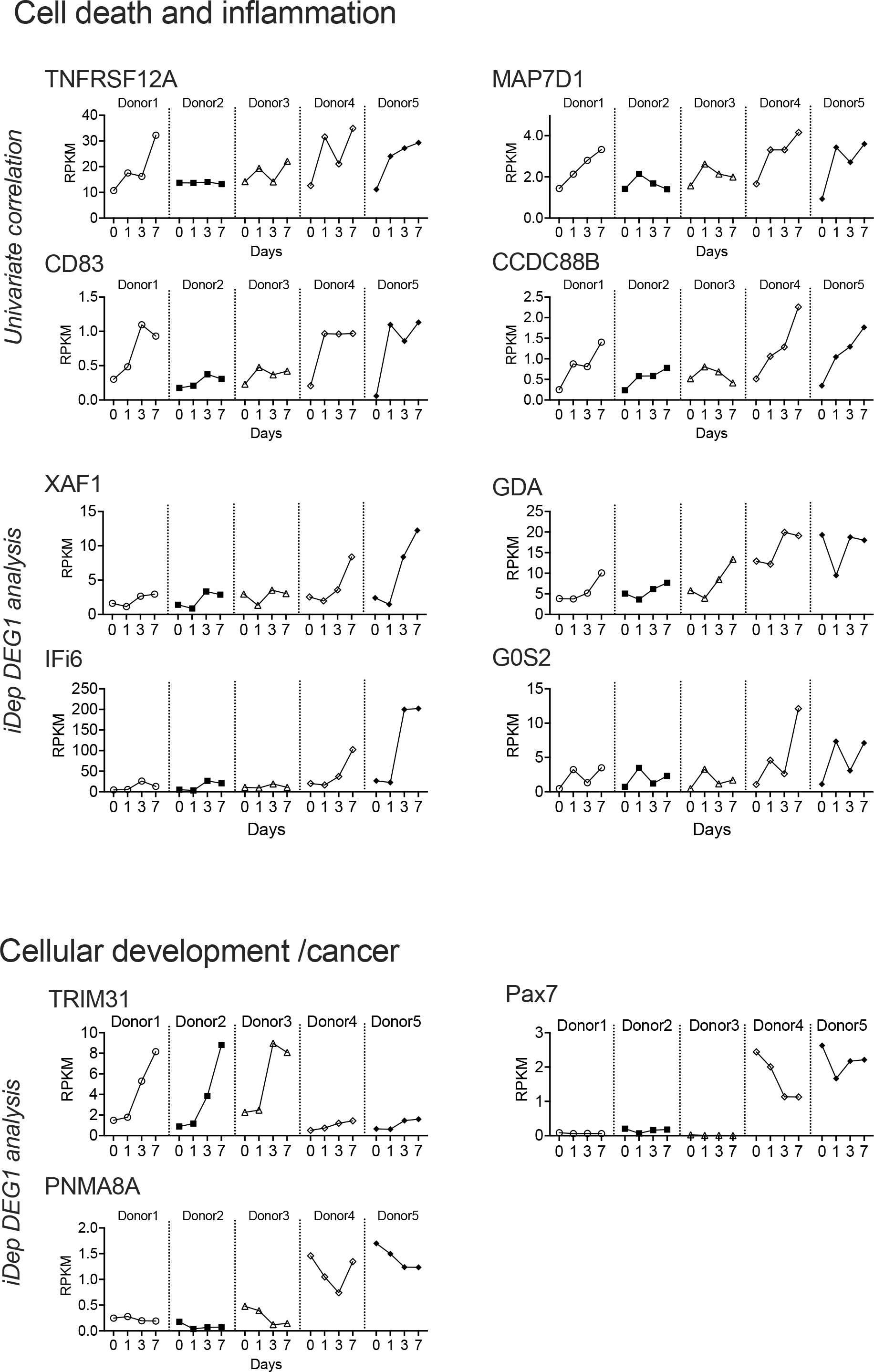
DEGs associated with NTHi susceptible donors 4 & 5. RPKM patterns are shown for each donor from day0 - day7. DEGs were identified using either univariate Pearson’s correlation or the i*DEP* DEG1 tool. For the univariate correlation analysis, only genes with a max RPKM > 1 are shown. All genes had absolute Pearson coefficient values of at least 0.75.

The i*DEP* analysis identified Pax7, G0S2, XAF1, PNMA8A, TRIM31, GDA and IFI6 as differentially expressed in donors 4 & 5 compared to donors 1-3 (fold-changes between 2.01 and 9.52 (IFI6); IFI6 & XAF1: adj.p-value < 0.1, others: adj.p-value<0.05)) (Figures 6, S7). Except for TRIM31, these genes were expressed at higher levels in donors 4 & 5, and interestingly, they appeared to fall into three functional groupings that partially overlap with the groups identified using univariate correlation. XAF1, TRIM31, G0S2 and IFI6 have known roles in apoptosis or its prevention, while PAX7, TRIM31 and PNMA8A are involved in cell development/development of cancer. Lastly, GDA has been proposed to affect microtubule assembly (60-65). Particularly striking differences in expression between the two donor groups were observed for PAX7, which lacked expression in donors 1-3, and XAF1, which showed a substantial increase in expression on days 3 and 7 in Donors 4 & 5 (Figures 6, S7).

Except for an accumulation of genes with known roles in apoptosis and inflammation, there was no overlap between the gene sets identified using our two approaches. This may, in part, be due to the relatively small number of donors, but also to the inherent differences in the two approaches taken. Interestingly, in some cases, particularly for donor 1, gene expression patterns were intermediate between those observed for donors 2 & 3 and donors 4 & 5 (e.g. G0S2, CD83), indicating a certain fluidity in gene expression patterns.

## Discussion

A characteristic property of *H. influenzae* is their persistence both as nasopharyngeal commensals and during upper and lower respiratory tract infections ^1, 2, 7, 46–48^. Here, we have shown for the first time that *H. influenzae* are effective long-term colonizers of primary human nasal epithelia (NHNE) with significant extra- and intracellular bacterial populations developing in all donors within 24 h p.i., and infections persisting for at least 7-14 days. The infection only minimally impacted NHNE physiology compared to uninfected controls and is consistent with an infection caused by a bacterium that is completely adapted to survival in contact with human respiratory epithelia as its only natural niche.

Underlying the apparently ‘normal’ physiology of the infected NHNE, however, we identified a highly dynamic response to the NTHi infection that included a strong hypoxia and an apparent inflammatory response in the first 24 h p.i. (Figures 1 & 7). However, this initial inflammatory response did not result in significant production of pro-inflammatory cytokines and bacterial clearance. Instead, gene expression patterns returned to an almost-pre-infection state by day3 p.i., while IL-8 and TNFα production was only observed on day7 and day14 p.i. (Figures 1, 5 & 7). We propose that these expression patterns indicate the development of NHNE tolerance to NTHi infection, a concept that is emerging in host-pathogen interaction studies ^49^. NHNE responses to NTHi infection in the first 72 h of our experiment and development of tolerance were highly similar in all five donors, suggesting a common, underlying mechanism.

**Figure 7.**
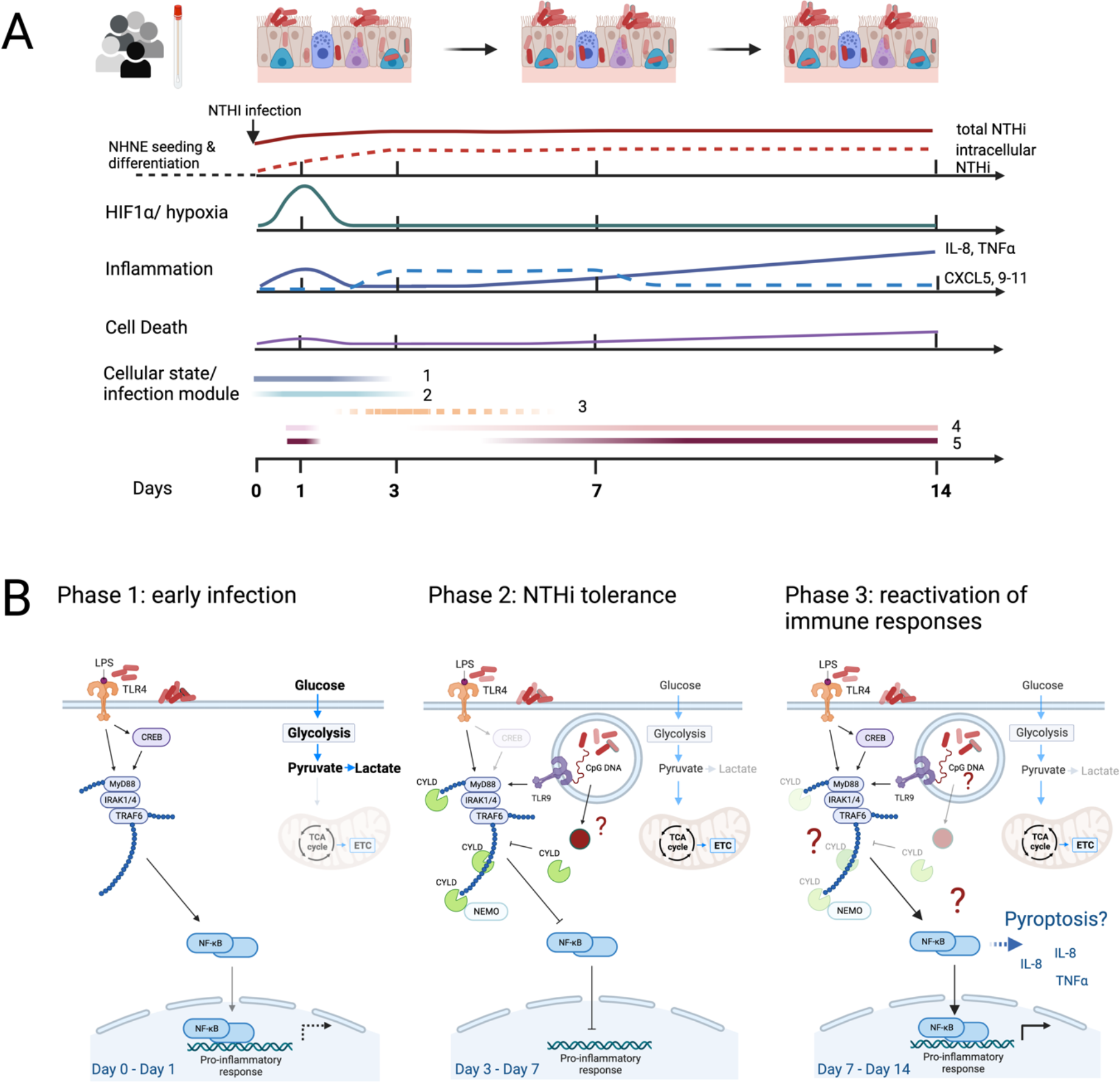
Schematic representation of NHNE responses to NTHi infection. **Panel A:** ‘Cellular state/ infection modules’ refers to infection modules identified by Avital et al. 2022 for macrophages. Module three was less enriched than the other modules in our sample and is therefore represented by a dashed line. **Panel B:** Model immunomodulatory effects of *H. influenzae* during infection of human epithelia. The figure was prepared using BioRender.com.

The observed epithelial tolerance of high bacterial burdens explains the known persistence of NTHi infections in humans, and further work will be needed to determine the contribution of molecular interactions between NTHi and NHNE to this process. NTHi intracellular populations are likely to play an active role in this previously undescribed effect as their development coincided with increasing NTHi ‘tolerance’ in NHNE. Molecular mechanisms involved in switching off the NHNE immune response could involve the CYLD deubiquitinase that has been shown to reverse NTHi-induced NFkB activation ^50, 51^. The reversal of the initial immune response and resulting tolerance to NTHi is also reminiscent of LPS insensitivity that has been described in macrophages^52, 53^. LPS insensitivity is a reversible process that results from repeated exposure to LPS and mainly affects pro-inflammatory effectors, while molecules involved in tissue repair functions are not subject to this desensitization ^52, 53^.

Interactions between epithelia and NTHi have been previously reported for infection of NHBE from a single human donor with the NTHi176 isolate strain^23^. In that study, the NHBE showed severe signs of degradation by 72 p.i., the final timepoint, and significant increases in IL-8 and TNFα production at 24h and 72h p.i., which differs markedly from our results. The most significant NHBE gene expression changes were associated with pathogen recognition, cellular junctions and cytoskeleton formation, ECM remodelling and inflammation^23^. Comparative analyses revealed that our data did not replicate the strong downregulation of genes involved in cytoskeleton and tight junction formation reported by^23^. However, genes involved in ECM remodelling and inflammation had similar overall up- and down-regulation trends in both studies, with the difference that in our study the relevant expression changes only occurred from day3 p.i., i.e. later than in ^23^.

Some of these differences may be explained by the differences in sampling times and possibly the use of primary bronchial rather than nasal cells in ^23^. We also cannot exclude that differences in the virulence of the NTHi strains used affected the host-pathogen interactions. However, all NTHi wild-type strains (n=3) we tested in preliminary infection experiments (data not shown &^19^) gave rise to stable NHNE infections, similar to what we reported in ^19^.

However, the similarity in expression trends in some gene groups and differences in the timing of these expression changes suggest that a donor specific temporal component may be common to NTHi infections in NHN/BE.

This type of temporal component was also apparent in the duration of the ‘NTHi-tolerance’ stage described above, which showed donor-specific differences. While all donors established NTHi tolerance by day3 p.i., some donors, i.e. donors 4&5, appeared to exit this tolerance stage faster than NHNE from other donors. This was indicated by an earlier onset of production of pro-inflammatory molecules, as well as increased cell death apparent by day14. This might be linked to several genes with differential expression patterns between donors 1-3 and the two less NTHi-tolerant donors. Most of these were linked to inflammation and cell death responses, but also restructuring of the microtubule network and cell proliferation, implicating these host processes in tolerance of NTHi and possibly also other bacterial infections.

Interestingly, if the analysis was extended to include day14 data, the altered gene expression trends observed in donors 4&5 by day7 p.i for, e.g., genes encoding TNFα, IL1β, IL-8, CXCL5, PLAU and G0S2 (Figures 6, S6 & S7) were also apparent in data for the ‘NTHi-tolerant’ donors 1-3, further suggesting a temporal component in the ‘exit’ of NHNE from the NTHi-tolerance phase (Figure S8). We propose that NTHi-infected NHNE from all donors followed a similar developmental trajectory, suggesting that NHNE infection with NTHi might proceed via distinct, conserved cellular states or ‘infection modules’. The concept of ‘infection stages’ or ‘modules’ was recently proposed for populations of macrophages, where infections with six different bacterial pathogens elicited responses corresponding to 5 conserved modules/ cellular states that were accessed to different extents depending on the infection scenario^54^. Interestingly, in our data, marker genes identified by ^54^ for these five modules showed hierarchical clustering that matched the NHNE infection status. Genes from modules 1-2 (‘uninfected cells’) were primarily associated with uninfected day 0 NHNE samples, while genes from modules 4 & 5 (‘high & very high inflammation’) were more likely to be highly expressed in day14 and day7 samples from our study where stronger inflammation phenotypes were observed (Figure S9). The relative activation of different ‘modules’ also matched the level of inflammatory responses seen in each donor, where samples from donor 2, that showed the least intense response to NTHi infection (Figures 6, S6, S7), clustered together with uninfected day0 samples from the other four donors (Figure S9).

Interestingly, analyses using only module 4 or module 5 genes in both cases revealed small clusters of active genes in samples collected on day1 p.i.. These clusters were distinct from genes that were strongly activated on days7 and 14 p.i. (Figure S9). This matches our observation that in NTHi-NHNE infections, a limited inflammation signature exists on day1 p.i., that is followed by a more robust and wide-ranging response to infection on days7 and 14 p.i.. Based on this analysis, we propose that responses to infection in human epithelial cells may be modular and resemble processes identified in canonical immune cells. Further work is needed to fully understand the responses of NHNE to NTHi infection, especially as our samples were derived from epithelia containing multiple different cell types and thus represent a mixture of cellular states, whereas the analysis of macrophage gene expression used single cell types and single-cell transcriptomes.

In summary, our data provide first evidence that NTHi infections can delay strong inflammatory responses in human epithelia and induce an apparent tolerance of NTHi infection that had not been previously observed, but could be a driver of NTHi persistence in the human respiratory tract. This state of ‘peaceful’ coexistence of NHNE and NTHi had a donor-specific component documented by longer or shorter delays in the reactivation of pro-inflammatory responses. This process is reminiscent of the LPS-insensitivity phenotype of macrophages, where differences in the duration and magnitude of insensitivity have also been observed in macrophages derived from different, common strains of wt mice such as C57Black or Balb/c^52, 53^. Our data also suggest that the concept of modular infection responses in human immune cells can likely be expanded to epithelial cells, that are the first line of defence in contact with many pathogens (Figure 7A). Whether bacterial effector proteins are involved in triggering NHNE tolerance of NTHi infection, and which activities within the human epithelia cause differences in tolerance of NTHi infections warrants further investigation.

## Acknowledgements

This research was supported by NHMRC grants 1158451 and 2019058 to UK and seed funding from the Australian Centre for Infectious Diseases (AID) to UK and PDS. Anna Henningham was the recipient of a UQ Postdoctoral Fellowship. PDS is a Leadership Fellow (L3) of the NHMRC.

## Author contributions

Data collection: UK, AH, MN, AB; Data analysis: AH, UK, EF, TS; Manuscript drafting: UK; visualization: UK, Manuscript editing: all authors

## Materials and methods

### Air liquid interface (ALI) differentiated normal human nasal epithelia (NHNE)

NHNE were prepared using primary nasal epithelial cells from five healthy (non-asthmatic and non-atopic) donors (ref. numbers: AV012, AV026, AV043, AV050 and AV058, Age 21-36 years old, designated as donors 1-5 or D1-D5 throughout). Nasal epithelial cells were harvested from the inferior surface of the anterior turbinate of consenting donors (UQ Ethics approval #2017000520) by curettage (curette: ASI Rhino-Prom Arlington Scientific, USA). Cells were resuspended in 2 ml of RPMI1640 medium and transported to the laboratory, where they were grown in submerged culture for approximately 3 weeks (passage 2) and then cryopreserved. For differentiation of primary nasal cells into NHNE, cells were seeded at an initial density of 25000 cells/transwell insert and cultured in submerged culture using Pneumacult-EX medium (Stemcell Technologies) until confluence was reached. Cells were airlifted and cultivated into well-differentiated epithelia as indicated by the presence of beating cilia under light microscopy using Pneumacult-ALI medium (Stemcell Technologies) in the basal chamber only. NTHi infections were carried out 27-35 days post-airlift. Transepithelial electrical resistance (TEER) measurements were carried out essentially as in ^55^, using an EVOM2 epithelial voltohmeter (World Precision Instruments) equipped with STX2 electrodes. For the measurements, 100 μL 1xPBS was added to the apical chamber of fully differentiated NHNE, and a mean resistance was calculated from triplicate measurements. Values were corrected for surface area and fluid resistance (insert without cells).

### Bacterial strain and culture conditions

Experiments used the non-typeable *H. influenzae* strain 86-028NP, an otitis media isolate ^56^. NTHi was routinely grown on Brain Heart Infusion agar (Becton Dickinson) supplemented with 10 μg/ml each of NAD and hemin and incubated at 37°C in the presence of 5% CO_2_.

### Infection of NHNE with non-typeable *Haemophilus influenzae*

NTHi were grown overnight on freshly prepared BHI agar plates, resuspended in 1x PBS to an initial OD_600_ of 0.15 and used to infect the NHNE apical side at MOI 10:1. Uninfected controls were treated with the same volume of 1xPBS. Infection was allowed to proceed for 24 h before non-adherent bacteria were removed using a surface wash with 100 μl 1x PBS. Routine apical washes (100 μl 1x PBS) and changes of the basal medium were carried out every 2-3 days. Wash fractions and media were collected and stored at −80°C. At each time point, total bacterial cell numbers were determined by lysing the NHNE using 100 μL of 1% saponin, followed by serial dilution of the lysate and plating on BHI agar plates to determine bacterial CFU/ml. To determine intracellular bacterial loads, NHNE were treated for 1h with 100 μL of gentamicin (100 μg/mL), followed by cell lysis and dilution plating.

### Immunofluorescence Microscopy

NHNE wells for microscopy were fixed by treatment with 3% paraformaldehyde, followed by washes with 1 × PBS and permeabilization using 0.5% Triton X-100. Blocking used 2% BSA, 0.2% Triton X-100 and was followed by incubations with the primary and secondary antibodies, essentially as in ^57^. Immunofluorescence staining used Phalloidin (F-Actin polymers, Thermo Fisher), TUNEL detection Kit (TUNEL488 probe, Roche) & DAPI (Sigma). The Anti-NTHi 86-028NP OMP antibodies ^58^ were a kind gift from Prof. Lauren Bakaletz.

### ELISA assays

Apical wash fractions collected pre- and post-infection were used to detect Lactate dehydrogenase activity and IL-8 concentrations using human LDH and IL-8 Alphalisa Kits (Perkin Elmer). ELISA Kits for the detection of human TNFα, EDN1 and LCN2 (NGAL) were purchased from ThermoFisher.

### Western Blots

Basal media samples collected during the infection process were separated on Bolt^TM^ 4-12% Bis-Tris gradient gels (ThermoFisher) using 20 μg protein/lane and Bolt^TM^ Loading Dye (ThermoFisher). Proteins were transferred to a nitrocellulose membrane using Power Blotter Select Transfer stacks and a Power Blotter (both ThermoFisher). Membranes were briefly stained with 1% PonceauS (w/v) to visualize protein, followed by destaining using 25 mM NaOH and blocking using 1x Blocker^TM^ FL Fluorescent protein block (ThermoFisher). Primary antibodies used were Anti-GapDH (G9545, Sigma-Aldrich, 1:2000), NGAL (PA5115502, ThermoFisher, 1:500), IL-8 (PA579113, Thermo Fisher, 1:500), TNFα (AMC3012, ThermoFisher, 1:1000), an Anti-Rabbit–Alexa conjugate antibody (A27041, ThermoFisher, 1:10 000) was used as the secondary antibody. All wash-steps between antibody incubations used 1xPBS, and membranes were imaged immediately after development using an Odyssey CLX imager (LiCor).

### RNA isolation and transcriptome analysis

RNA was isolated from infected and uninfected NHNE using TRIZOL reagent (Life Technologies). For each sample, membranes from 3 replicate transwells were excised using a scalpel blade and immediately immersed in 200 μL of TRIZOL. Epithelial cell layers were then removed from the membrane by gentle pipetting. For bacterial RNA isolation, cell material from the same BHI agar plate used to prepare the inoculum was collected using a sterile inoculation loop and resuspended in 200 μL TRIZOL. Samples were frozen at −80° C, and RNA isolated as per the manufacturer’s instructions within 2 weeks of sample collection. No RNA could be obtained for Donors 4 & 5 samples from day14 due to epithelial damage. All RNA samples were treated with EZ-DNAse (Life Technologies) to remove genomic DNA, followed by clean-up using the RNA Clean & Concentrate Kit-5 (ZYMO). RNA Integrity was determined using RNA screen tape (Agilent) and a Tapestation 4200 (Agilent). Samples with an RNA Integrity Number (RIN) of 8 or more were used for rRNA depletion using a Ribominus v2 kit (ThermoFisher) and both bacteria-specific and human-specific rRNA probes as per the manufacturer’s instructions (Life Technologies). Ribo-depleted RNA was concentrated using the RNA Clean & Concentrate Kit-5 (ZYMO), and integrity and concentrations of the ribo-depleted RNA checked using a High sensitivity RNA Tape (Agilent) and Qubit RNA assay (Life Technologies). Pair-ended libraries (2×100 bp, TruSeq) were prepared from ribo-depleted RNA and separated on a NovaSeq 6000 Flowcell at the Ramaciotti Centre for Genomics (University of Sydney). Data analysis used the CLC Genomics workbench (Qiagen, v.21) ‘RNAseq and Differential Gene Expression Analysis’ workflow. Data were mapped to the NTHi 86-028NP (NCBI acc.no: NC_007146.2) or the human (hg38) genome or transcript data. Any remaining rRNA reads were removed prior to statistical analyses. The RNA-seq raw data was high quality, with average FastQC Phred scores between 37 and 40. Approximately ∼ 70 million reads were collected per sample, of which about 65 million RNA reads mapped to the human genome, giving approximate genome coverages of 100 times for NTHi and ∼4.5 times for the human samples. One NTHi sample (D2 day3) was removed from the analysis as it had less than 2 million reads and behaved as an outlier in all analyses. Differentially expressed genes were classified as showing at least 2-fold up-/down-regulation and either an FDR p-value <0.05 (bacterial genes) or a Bonferroni corrected p-value p<0.05 (human genes).

Analyses to identify relevant regulators and pathways in NHNE data used Ingenuity Pathway Analysis (IPA, Qiagen). In IPA canonical pathway analyses, a z-score above 2.0 was deemed significant, and only datasets with samples for all five donors, i.e. days 0, 1, 3 and 7 were evaluated to enable robust statistical analyses. Gene Ontology (GO) enrichments used the geneontology web tool (www.genetontology.org) that is based on Panther GO for NTHi DEGs ^59^ and the Toppfun tool for NHNE GO enrichments ^60^. BlastKOALA was used to assign KEGG orthology categories to proteins encoded by DEGs for NTHi and NHNE ^61^. The CLICK algorithm embedded in the EXPANDER suite of programs was used to identify common gene expression patterns using default settings and RPKM values as the input ^62, 63^.

### Analysis of gene expression changes between NHNE

Univariate Pearson’s correlation was used to look for differentially expressed genes that were common among donors 1-3 on all days and donors 4-5 only on day0 (non-severe class) and different to donors 4-5 on days 1-3-7 (severe class). By taking donors 4-5 on day0 into the same response variable class as donors 1-3 on all days (non-severe class), the analysis focused on detecting differentially expressed genes that are donors 4-5 specific following exposure to the pathogen. Additional comparisons between donor groups (donors 1-3 and donors 4 & 5) were performed using the *iDEP* platform DEG tools ^64^.

### Sequence data availability

Transcriptome sequencing data have been deposited with NCBI as bio project PRNJ94815 and will be made available on publication of the manuscript. Visualizations used GraphPad Prism v9.4, GraphBio ^65^ and BioRender.com.

## Supplementary Information

Supplementary Figures – Figures S1-S9, word document

Supplementary Table 1 – NTHi DEGs and GO term enrichments

Supplementary Table 2 – NHNE DEGs

Supplementary Table 3 – NHNE GO term enrichments TOPPFUN

Supplementary Table 4 – NHNE IPA analyses

Supplementary Table 5 – NHNE donor group (D1-3 vs D4-5) analyses

