## Supplementary Figures for "Tolerance to *Haemophilus influenzae* infection in human epithelial cells: insights from a primary cell-based model"

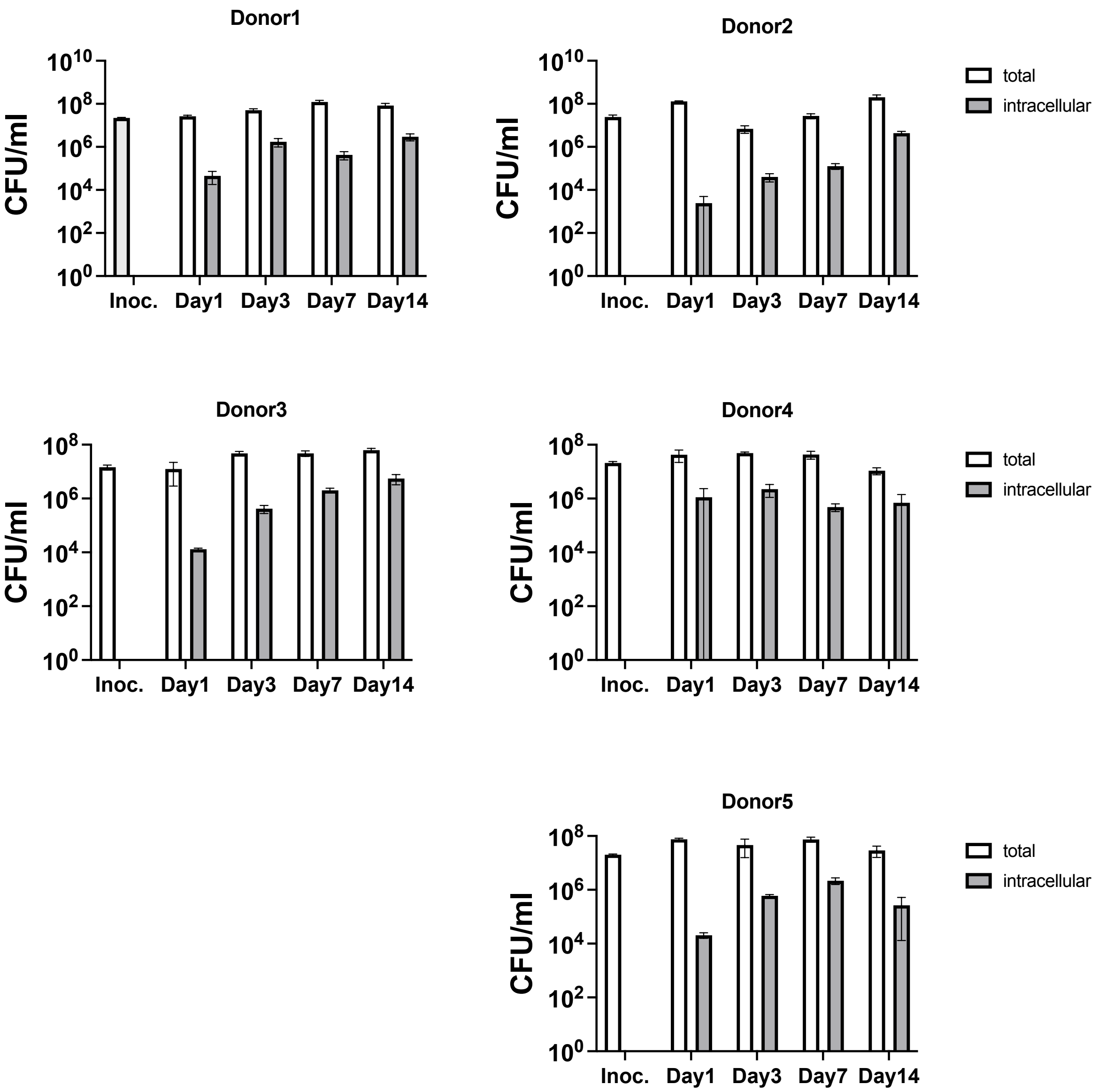


**Figure S1** – Bacterial Loads in in NHNE from different donors. Inoc. - inoculum

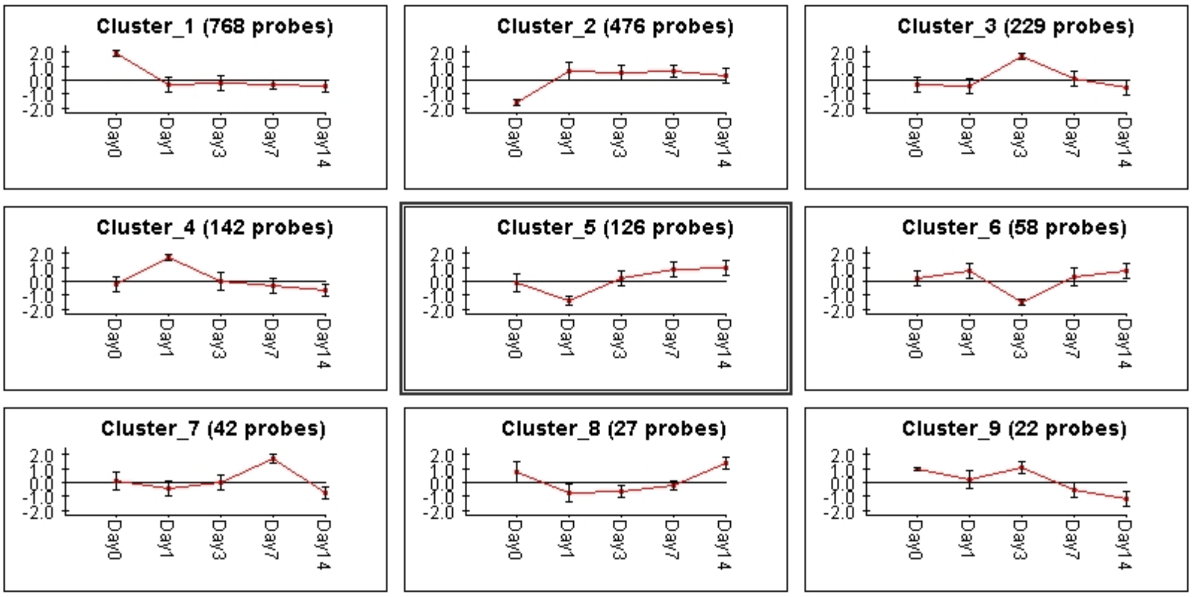


**Figure S2** – Gene expression clusters (fold-canges) identified for NTHi using the CLICK algorithm integrated into the EXPANDER package. Using default settings, nine clusters were identified, with only 44 genes not mapped. Clusters 1,5,7 & 9 show a decrease in gene expression from day 0 to Day1. Genes in Cluster 3 also showed this feature but to a lesser degree. Cluster homogeneity values were between 0.803 and 0.883.

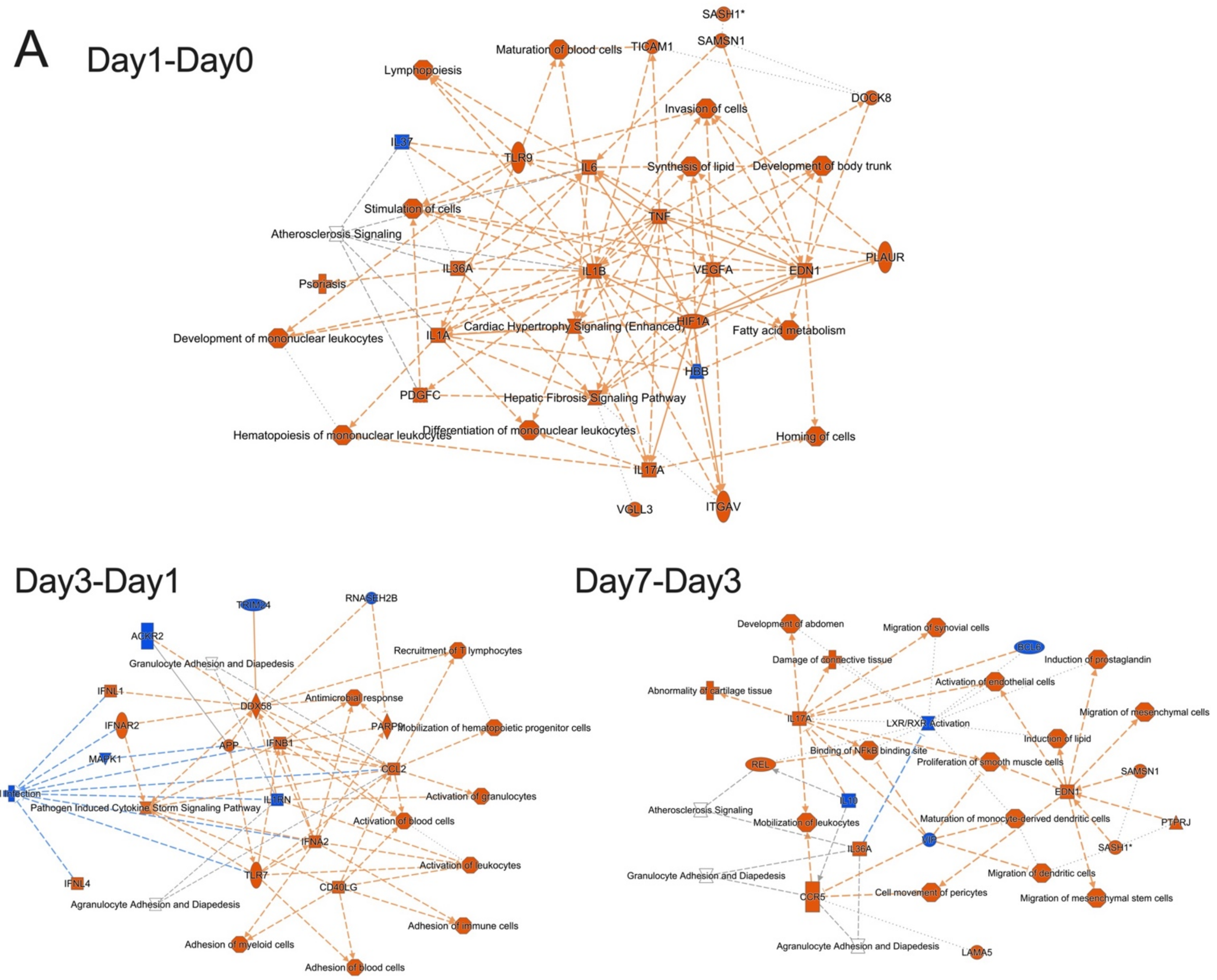
A
Day1-Day0
SASH1*
SAMSN1
Maturation of blood cells
TICAM1
Lymphopoiesis
DOCK8
Invasion of cells
IL37
TLR9
IL6
Synthesis of lipid
Development of body trunk
Stimulation of cells
TNF
Atherosclerosis Signaling
IL36A
IL1B
VEGFA
EDN1
PLAUR
Psoriasis
Development of mononuclear leukocytes
IL1A
Cardiac Hypertrophy Signaling (Enhanced)
HIF1A
Fatty acid metabolism
HBB
PDGFC
Hepatic Fibrosis Signaling Pathway
Hematopoiesis of mononuclear leukocytes
Differentiation of mononuclear leukocytes
Homing of cells
IL17A
VGLL3
ITGAV
Day3-Day1
TRIM24
RNASEH2B
ACKR2
Granulocyte Adhesion and Diapedesis
Recruitment of T lymphocytes
IFNL1
Antimicrobial response
DDX58
IFNAR2
PARP9
Mobilization of hematopoietic progenitor cells
APP
IFNB1
MAPK1
CCL2
Viral Infection
IL1RN
Activation of granulocytes
Pathogen Induced Cytokine Storm Signaling Pathway
Activation of blood cells
IFNA2
TLR7
Activation of leukocytes
IFNL4
CD40LG
Agranulocyte Adhesion and Diapedesis
Adhesion of immune cells
Adhesion of myeloid cells
Adhesion of blood cells
Day7-Day3
Development of abdomen
Migration of synovial cells
BCL6
Damage of connective tissue
Induction of prostaglandin
Activation of endothelial cells
Abnormality of cartilage tissue
Migration of mesenchymal cells
IL17A
LXR/RXR Activation
Induction of lipid
Binding of NFkB binding site
REL
Proliferation of smooth muscle cells
EDN1
SAMSN1
Atherosclerosis Signaling
Mobilization of leukocytes
Maturation of monocyte-derived dendritic cells
PTPRJ
IL36A
VIP
SASH1*
Granulocyte Adhesion and Diapedesis
Migration of dendritic cells
Cell movement of pericytes
Migration of mesenchymal stem cells
CCR5
Agranulocyte Adhesion and Diapedesis
LAMA5

B Day1-Day0

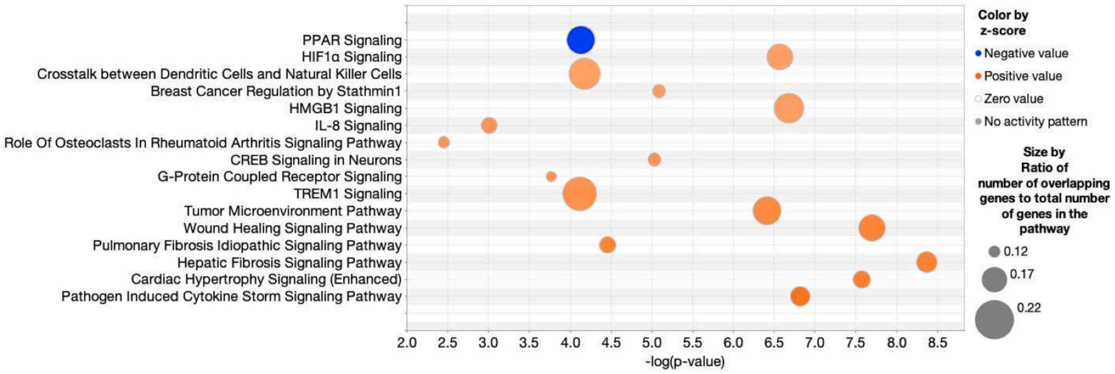


Day3-Day1

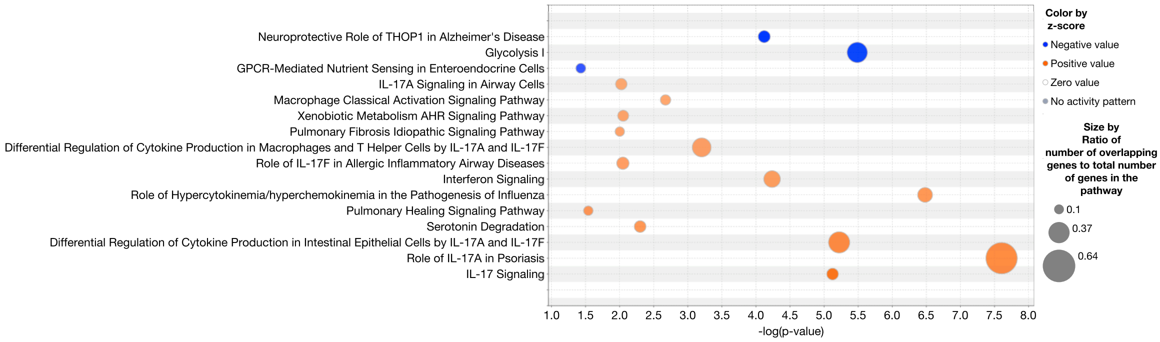


Day7-Day3

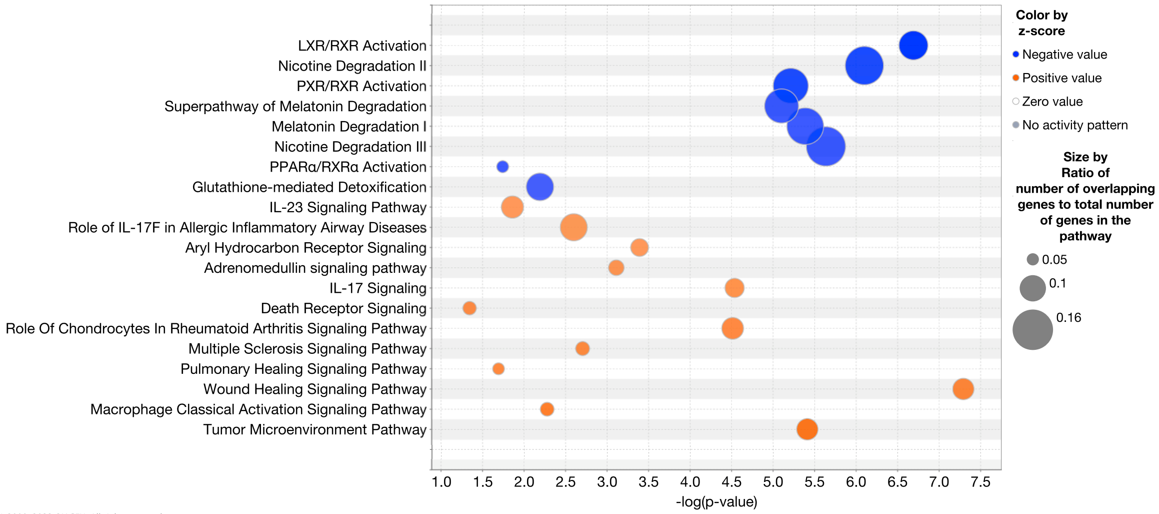

C

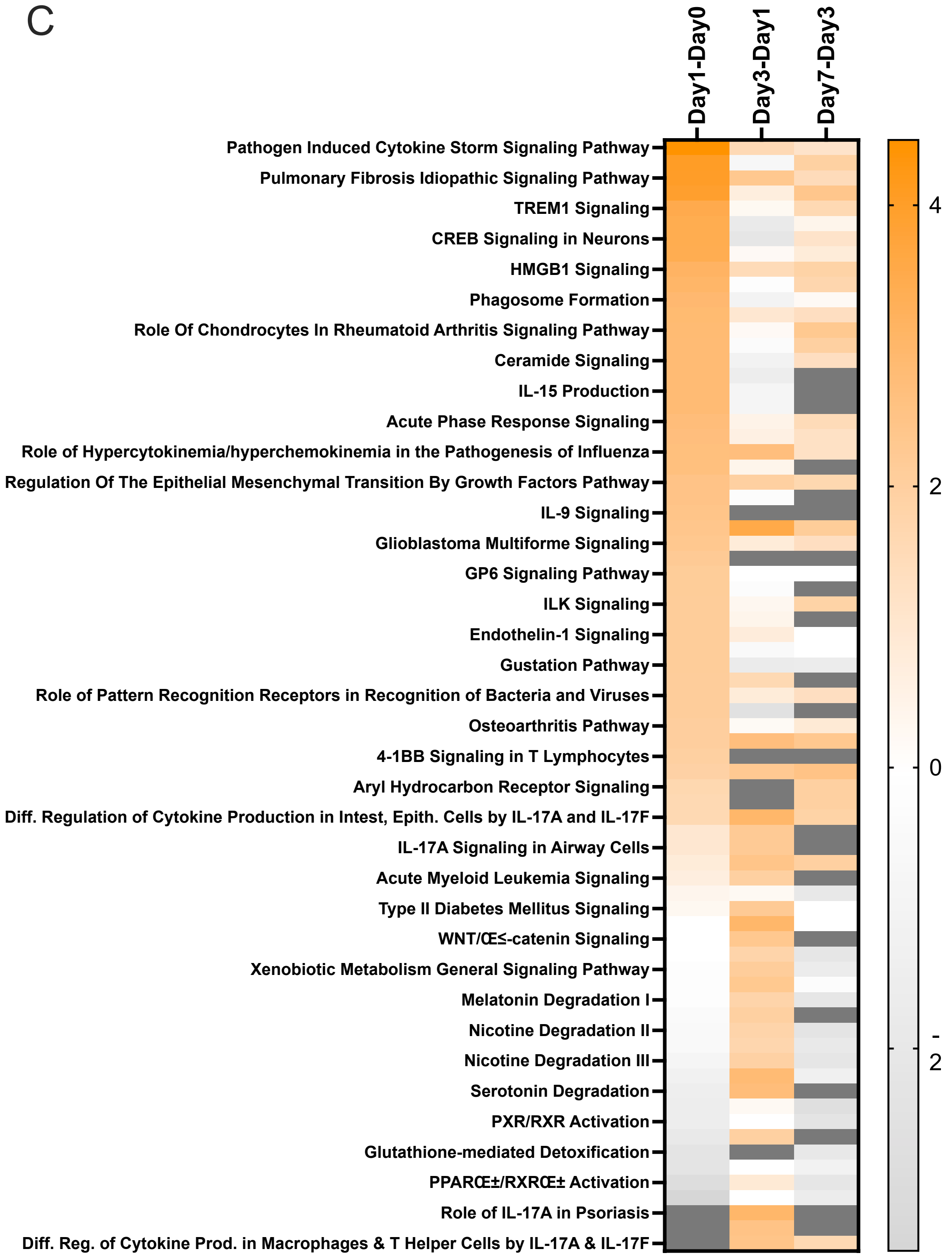

Day1-Day0
Day3-Day1
Day7-Day3
Pathogen Induced Cytokine Storm Signaling Pathway
Pulmonary Fibrosis Idiopathic Signaling Pathway
TREM1 Signaling
CREB Signaling in Neurons
HMGB1 Signaling
Phagosome Formation
Role Of Chondrocytes In Rheumatoid Arthritis Signaling Pathway
Ceramide Signaling
IL-15 Production
Acute Phase Response Signaling
Role of Hypercytokinemia/hyperchemokinemia in the Pathogenesis of Influenza
Regulation Of The Epithelial Mesenchymal Transition By Growth Factors Pathway
IL-9 Signaling
Glioblastoma Multiforme Signaling
GP6 Signaling Pathway
ILK Signaling
Endothelin-1 Signaling
Gustation Pathway
Role of Pattern Recognition Receptors in Recognition of Bacteria and Viruses
Osteoarthritis Pathway
4-1BB Signaling in T Lymphocytes
Aryl Hydrocarbon Receptor Signaling
Diff. Regulation of Cytokine Production in Intest, Epith. Cells by IL-17A and IL-17F
IL-17A Signaling in Airway Cells
Acute Myeloid Leukemia Signaling
Type II Diabetes Mellitus Signaling
WNT/Œ≤-catenin Signaling
Xenobiotic Metabolism General Signaling Pathway
Melatonin Degradation I
Nicotine Degradation II
Nicotine Degradation III
Serotonin Degradation
PXR/RXR Activation
Glutathione-mediated Detoxification
PPARŒ±/RXRŒ± Activation
Role of IL-17A in Psoriasis
Diff. Reg. of Cytokine Prod. in Macrophages & T Helper Cells by IL-17A & IL-17F
4
2
0
-
2

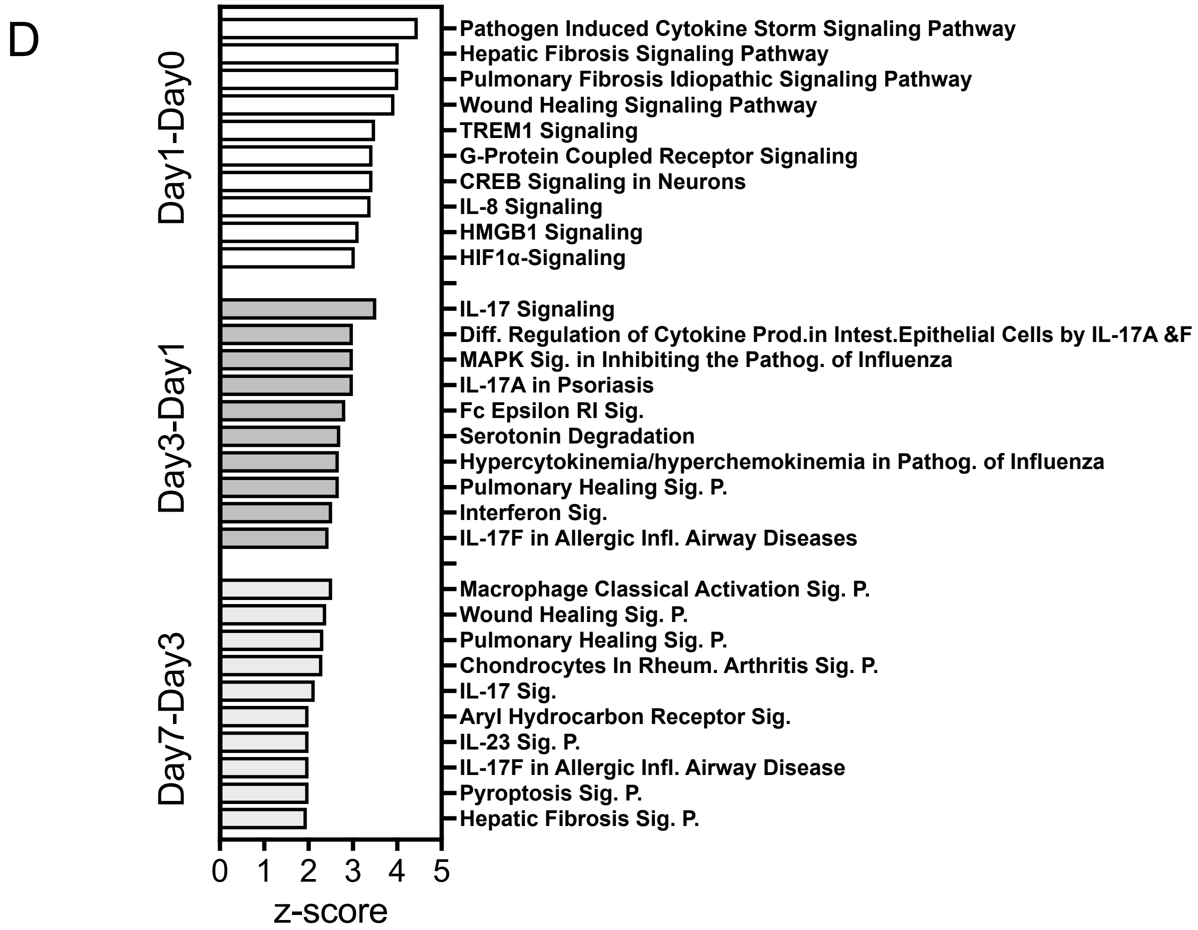


**Figure S3** – Figures from NHNE RNAseq data using Qiagen IPA. **Panel A**: Effector summary networks **Panel B**: Top Canonical Pathways identified in data comparisons displayed as bubble plots. Z-score cut offs: Day1-Day0: 3.0; Day3-Day1: 2.2, Day7-Day3: 2.0, FDR p-value <0.05. **Panel C:** Heatmap of canonical pathways with z-score of at least 2 in one of the three comparisons. Data is sorted in descending order for the Day1-Day0 data. Colour Dark gray- pathway not identified. **Panel D:** Top 10 Canonical Pathways and their z-scores

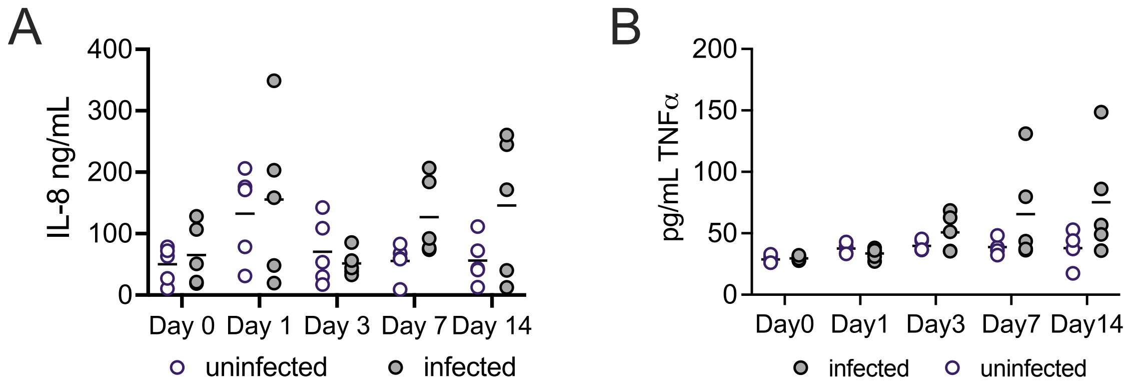


**Figure S4** – Detection of IL-8 (A) and TNFα (B) in NHNE apical wash fluid using ELISA. Each datapoint represents an average of at least n=3 replicate wells for each donor.

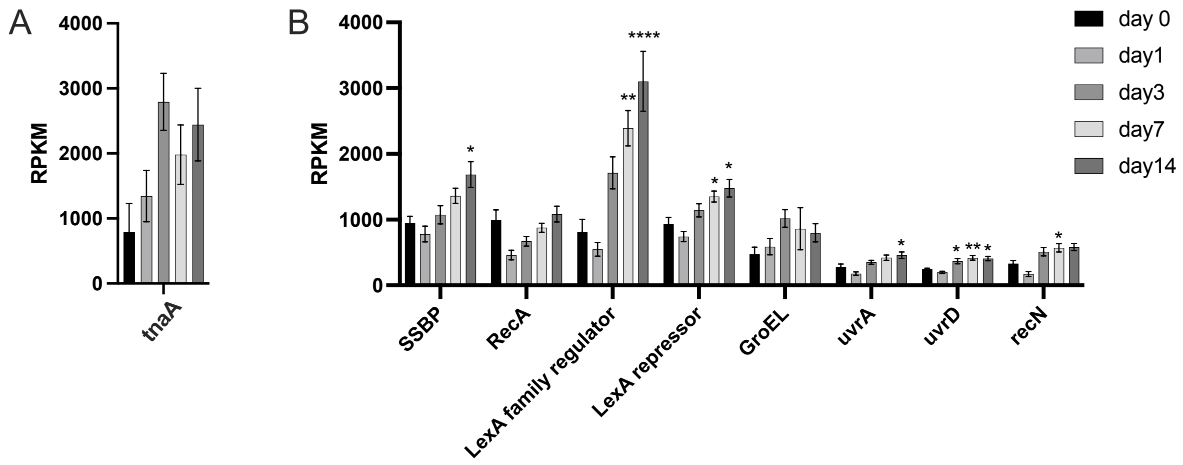


**Figure S5** – Expression of genes involved in NTHi stress responses including SOS responses. Panel **A**: tnaA tryptophanase, Panel **B**: genes involved in SOS responses. RPKM values shown are averages of values obtained from n=5 donors (Donors 1-5), except for the Day14 values (n=3, Donors 1-3). Statistical analyses used 1-Way ANOVA, using the Day0 value as the reference. p-values: *<0.05, **<0.01, **** <0.0001

### Cell death and inflammation

TNFRSF12A

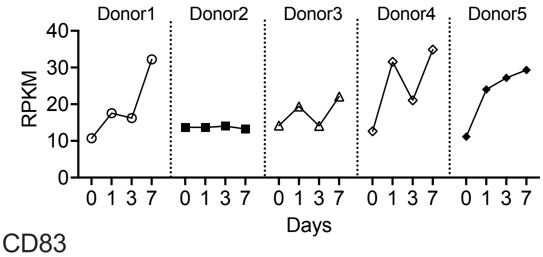


MAP7D1

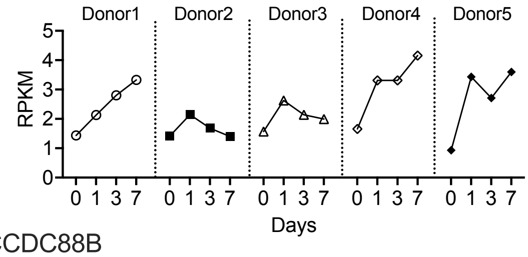


CD83

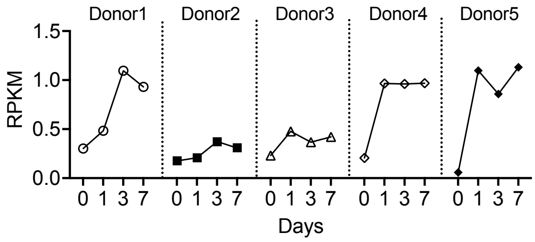


CCDC88B

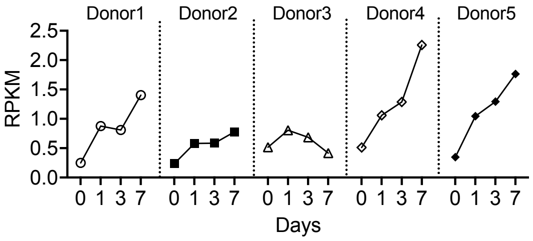


### Gene regulation

ZNF107

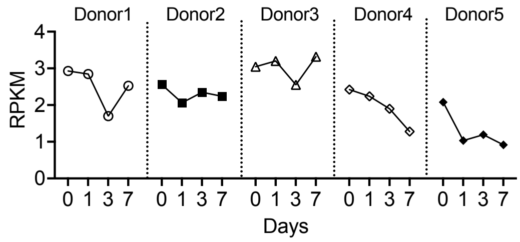


### Other functions: developmental and neuronal signalling

PHLDA2

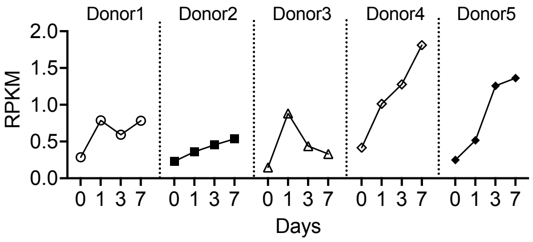


PHETA1

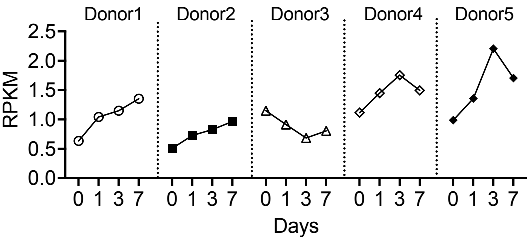


MRGPRX3

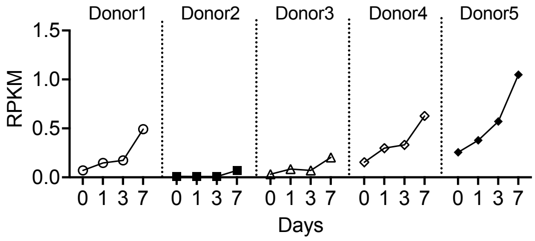


**Figure S6** – DEGS associated with NTHi susceptible donors 4 & 5 identified using univariate Pearson's correlation. RPKM patterns are shown for each donor from Day0 - Day7. Only genes with a max RPKM or > 1 are shown. All genes had Pearson coefficients with an absolute value of at least 0.75.

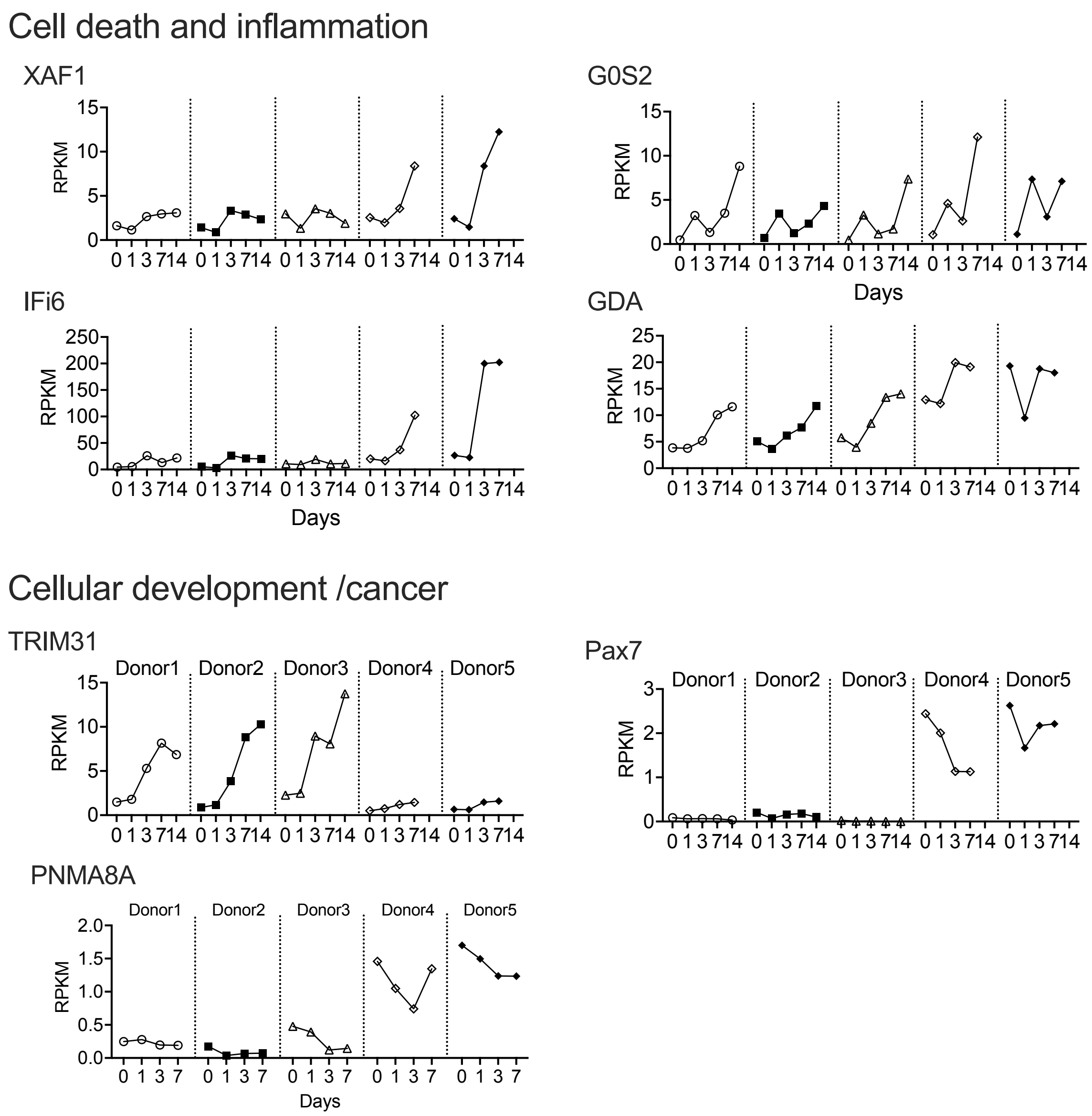


**Figure S7** - DEGS associated with NTHi susceptible donors 4 & 5 identified by idep.94. RPKM patterns are shown for each donor from Day0 - Day7 (Day14 for donors 1-3). DEGs were identified using the DEG1 tool within the idep.94 platform.

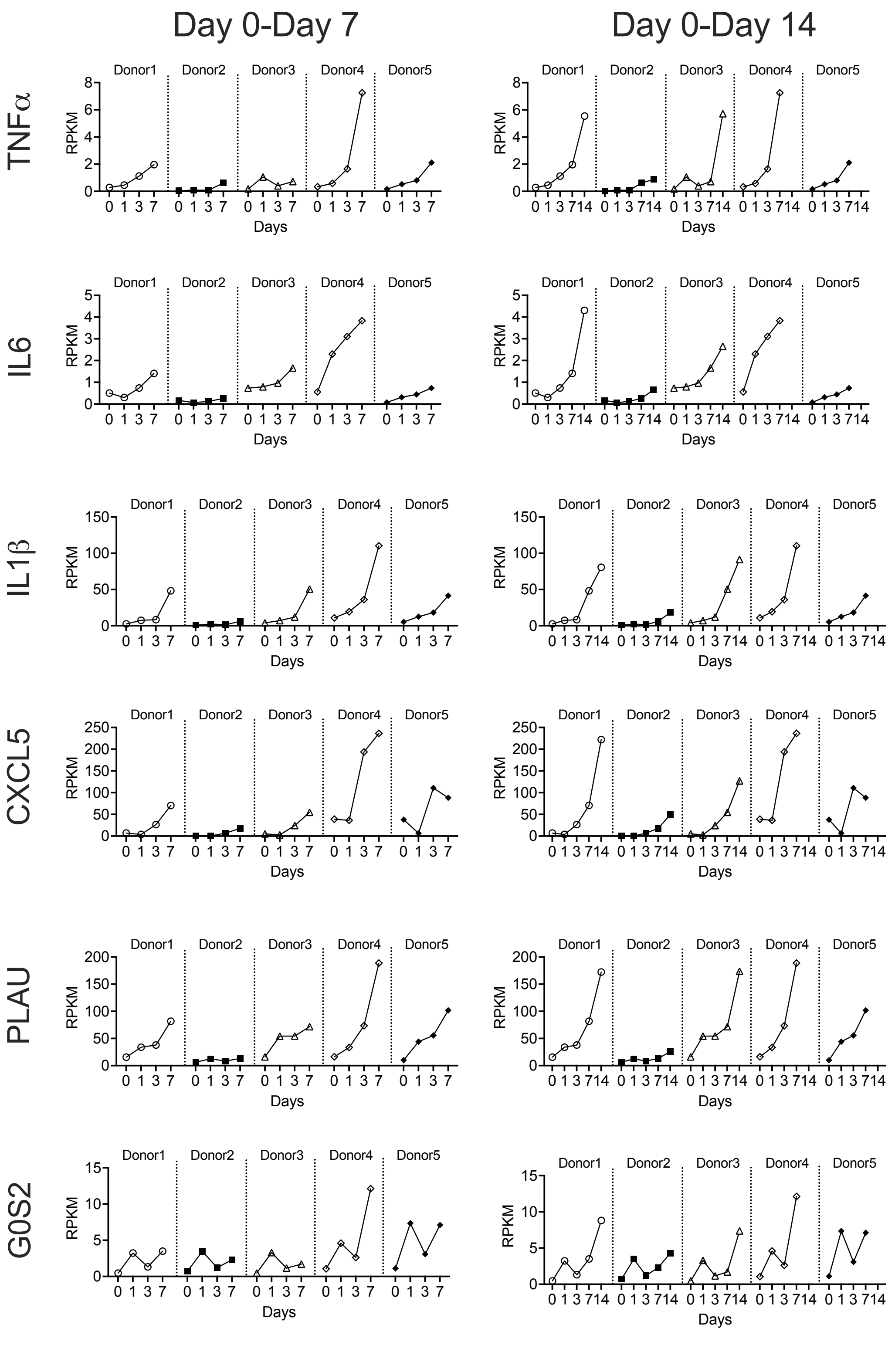


**Figure S8 Gene expression patterns for TNFα, IL6, IL1β, CXCL5, PLAU and G0S2 for all donors**. Left diagrams show expression values from Day0 to Day7, right diagrams include RPKM values for Day14 for donors 1-3, highlighting that when considering this extra datapoint, gene expression patterns across donors look more similar than in the Day0- Day7 comparison.

A

Module 1-5 genes

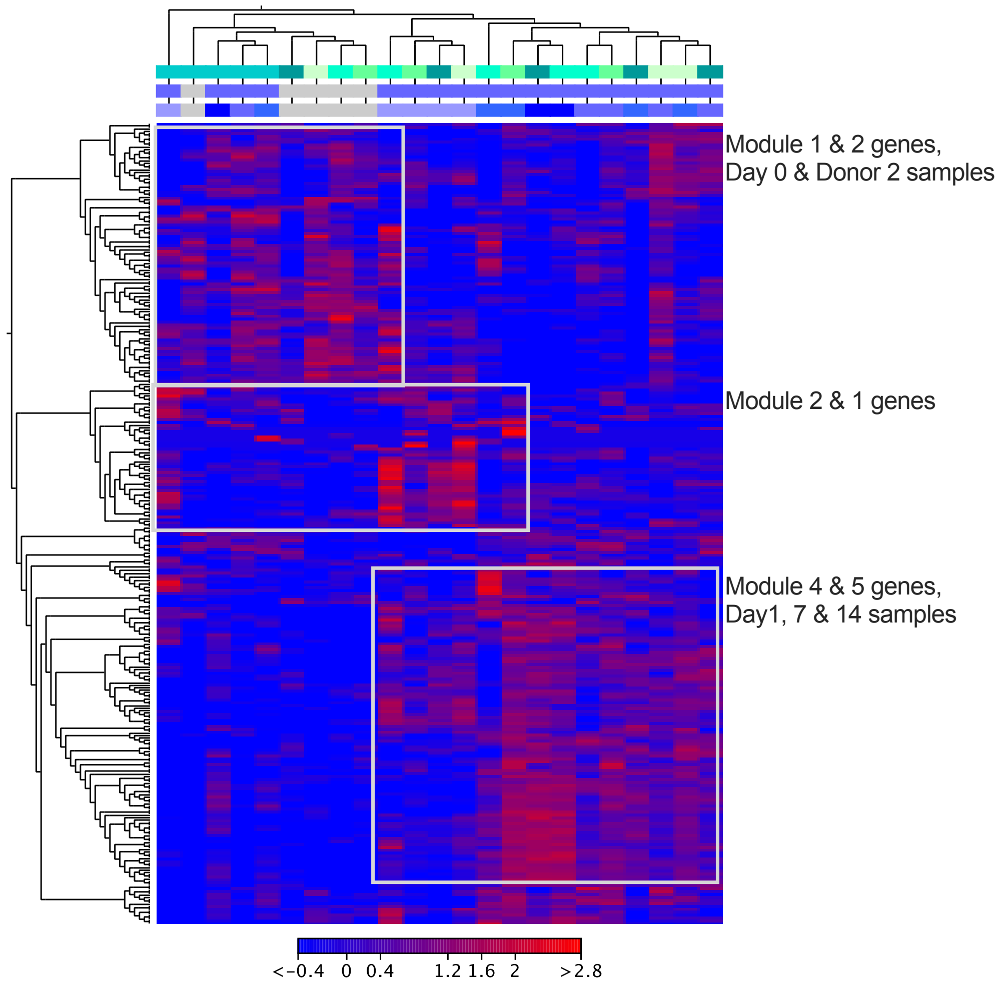

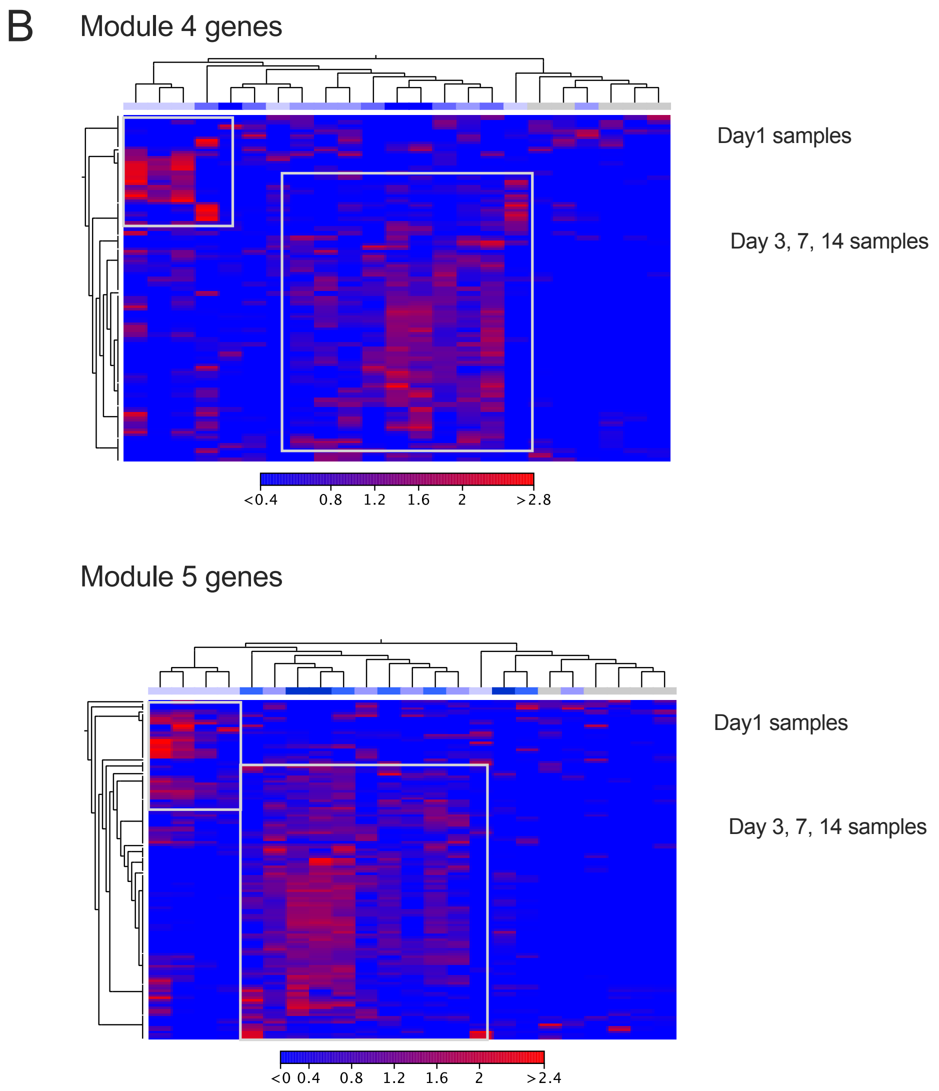


**Figure S9:** Heatmaps of infection module genes as defined by Avital et al 2022 for macrophages. **A:** Heatmap using genes for all five gene expression modules identified by Avital et al. **B:** Heatmap for module 4 and module 5 genes.

Colour bars (top of heatmaps): *Heatmap A:* top: green colours – donors, middle: grey – uninfected, blue – infected, bottom: sampling time, grey Day 0, blue colours: lightest shade – day 1, Darkest shade - day 14. *Heatmap B:* sampling time, grey Day 0, blue colours: lightest shade – day 1, Darkest shade - day 14
